# Biophysically relevant network model of the piriform cortex predicts odor frequency encoding using network mechanisms

**DOI:** 10.1101/2025.07.14.664633

**Authors:** Ankit Kumar, Om Chaudhari, Manan Jindal, Ashutosh Modi, Debanjan Dasgupta

**Affiliations:** Neural Circuit Dynamics Lab, Department of Biological Sciences and Bioengineering, Indian Institute of Technology, Kanpur, 208016, India; Department of Computer Science and Engineering, Indian Institute of Technology, Kanpur, 208016, India; Department of Electrical Engineering, Indian Institute of Technology, Kanpur, 208016, India

**Keywords:** Circuit motifs, odor frequency, olfaction, piriform cortex

## Abstract

Olfactory-guided animals utilize fast concentration fluctuations in turbulent odor plumes to perceive olfactory landscapes. Recent studies have demonstrated that the olfactory bulb (OB) encodes such temporal features present in natural odor stimuli. However, whether this temporal information is encoded in the piriform cortex (PCx) remains unknown. Hence, we developed a biophysically relevant PCx network model and simulated it using previously recorded *in vivo* activities of mitral and tufted cells in response to 2Hz and 20Hz odor frequencies for three stimulus mixtures. Analysis of single-cell activity across trials revealed that individual pyramidal neurons (PYRs) were largely ineffective at discriminating between 2Hz and 20Hz. However, the trial-averaged activity of the PYR population could discriminate between the two frequencies significantly. Moreover, using log-likelihood scores we further discovered that odor frequency discrimination happened through a highly distributed mechanism among the PYRs. One-dimensional convolutional neural network models trained and tested on PYRs’ activities achieved discrimination accuracies up to 95%. Using virtual synaptic knockout models, we found that eliminating either feedback or feedforward inhibition onto PYRs improved the decoding accuracy across all odor conditions. Conversely, eliminating recurrent excitation among PYRs or simultaneously eliminating recurrent inhibition within both interneuron populations degraded decoding performance. Removing recurrent connections within individual interneuron populations had minimal effects on the performance. Overall, our PCx model demonstrates that it can discriminate between 2Hz and 20Hz odor stimuli, with a bidirectional capability of performance modulation by specific circuit motifs. These findings predict that the piriform cortex encodes and processes temporal features of odor stimuli.

**New & Noteworthy:** We simulated the first biophysically relevant network model of the piriform cortex (PCx) to show differential encoding of odor frequencies at 2Hz and 20Hz. 1D convolutional neural networks demonstrated a distributed role of pyramidal neurons in the encoding. Surprisingly, eliminating feedforward or feedback inhibition improves frequency discrimination, while eliminating recurrency impairs it. Specific circuit motifs, not just baseline activity levels, determine odor frequency representation. Overall, our results predict the PCx’s capacity for temporal odor processing.

## Introduction

In the natural olfactory environment, odorant molecules travel as odor plumes due to the turbulence of natural airflow (1–4). These odor plumes are characterized by rapid concentration fluctuations of odorant molecules over space and time (4–8). It has been observed that such odor plumes carry crucial spatial information about odor source location, embedded in their temporal structure (1,3,7). Olfactory-guided animals can perceive the temporal structure of odors (1,3,7,9,10), and recent reports indicate that humans can perceive such stimuli as well (11). Early olfactory regions in the nervous system, like the antennal lobe in insects (12–17) and the olfactory bulb (OB) in mammals, can represent these temporal features (18,19). However, whether higher olfactory cortical areas such as the piriform cortex (PCx) encode this temporal information remains unknown.

Owing to the slow signal transduction kinetics of GPCR-based odorant receptors, olfaction has largely been thought to be devoid of high-frequency temporal features of odor stimuli (20). However, studies in invertebrates (15,16) and more recently in mice (3,8,10,18,19) and humans (11) have shown that different temporal features of odor stimuli can be perceived and encoded at least in the early olfactory system. One of the main cortical regions associated with olfaction, the PCx, has primarily been implicated in encoding odor identity and intensity (21,22). The recurrent network of the PCx is responsible for olfactory sensory processing, which includes shaping the ensembles of odor-identity encoding piriform neurons (23), gain control, pattern separation and pattern completion, as well as odor learning and memory formation (24–28). Moreover, PCx, through its cortical feedback axons, provides the OB with efficiently formatted and decorrelated information about diverse odorant mixtures and concentrations (29). However, we completely lack an understanding of whether PCx’s recurrent circuitry can encode odor temporal features such as frequency of odor concentration fluctuations. Hence, given the importance of odor temporal information in various ethological behaviors and the known involvement of the olfactory cortex (30,31), we investigated whether the PCx is capable of encoding different temporal features present in naturalistic odor stimuli.

To investigate this, we developed a biophysically constrained network model of the PCx. We analyzed pyramidal neurons (PYRs) activities in response to three odorant mixtures presented at 2 Hz and 20 Hz frequencies. Additionally, we simulated various PCx synaptic knockout models to determine the extent to which each circuit motif influences the network’s frequency discrimination capacity. Our findings demonstrate that the PCx network can effectively discriminate between 2 Hz and 20 Hz odor frequencies harnessing a network-wide distributed mechanism. Eliminating direct feedforward or feedback inhibition onto the PYRs enhanced frequency decoding capacity of the PCx network model, whereas removing recurrent excitation onto the PYRs or disinhibition within both feedforward and feedback interneurons impaired this capacity. In contrast, the odor frequency decoding accuracy was minimally affected by the synaptic disconnections of the intra-feedforward or intra-feedback interneurons. Overall, our study predicts that the PCx network can perform odor frequency discrimination using distributed network mechanisms. Furthermore, odor frequency encoding can be modulated bidirectionally by the different synaptic motifs within the network.

## Materials and Methods

### Model

The code was written in Python using the NEURON 8.0.2 simulation environment (32) and executed using Conda and Python virtual environments. The integration time step for the simulations was 0.025 ms.

### Model Architecture

The piriform cortex model consisted of three cell types: 500 excitatory pyramidal cells (PYRs), 50 feedforward inhibitory neurons (FFINs), and 50 feedback inhibitory neurons (FBINs). All synaptic connections within the network were randomly assigned, regardless of cell location. Previously recorded *in vivo* activities from 20–50 unique mitral/tufted cells (M/Ts) from the olfactory bulb in response to different stimulus conditions were used as input to simulate the PCx network model (18). These spiking activities were duplicated to form an M/T cell input pool of size equivalent to ∼22.5% of the total PYR population. Input to each pyramidal neuron was randomly selected from 20% (33) of this M/T cell pool. Each pyramidal neuron was connected to 20% of the other pyramidal neurons in the population (33) via proximal apical dendrites (23). The FFINs projected to the distal apical dendrites of PYRs (34–36). Each PYR was inhibited by 30% of the FFINs (36). Each FFIN received inputs from 40% of M/Ts (35,36). Each FBIN received excitatory input from 18% of PYRs, while each PYR received somatic inhibition from 35% of FBINs (36). Each FBIN received inhibitory inputs from 10% of other FBINs, while each FFIN received inhibitory inputs from 40% of other FFINs.

### Neuron and Electrophysiological parameters

#### Pyramidal (PYR) neuron model

Each PYR was composed of a soma and five dendritic sections. Both the length and diameter of the somatic section were 29 µm. The length of all dendritic sections was 100 µm. The diameters of all the dendritic sections 1–5 were 2.57, 1.77, 1.2, 0.8, and 0.5 µm, respectively (Fig. S1A). Here, dendritic section 5 was considered the distal apical dendrite. The charge balance equation for each neuronal compartment is described by:

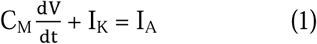

Where:

C_M_ is 1 µF/cm^2^

V is membrane voltage

I_K_ is the sum of ionic currents that pass through the membrane, and

I_A_ is the current entering from adjacent regions.

The equations governing I_K_ are defined explicitly in the mod file of each ion channel mechanism. Ion channel mechanisms (*.mod files) were compiled using NRNIVMODL in NEURON. The axial resistance (R_a_) for all neuronal compartments was set to 173 Ω·cm. Each PYR section had sodium and delayed rectifier potassium channel mechanisms adopted from Migliore et al. (1999). Leak conductance in PYR was implemented using NEURON’s built-in “leak” distributed mechanism. The maximum conductance of sodium, potassium, and leak ion channels was 0.5, 0.06, and 0.003 S/cm^2^, respectively. The reversal potential of sodium, potassium, and leak ion channels was 50 mV, −85 mV, and −75 mV, respectively. The resting membrane potential (RMP) of PYRs was −74.1 ± 8.6 mV (mean ± SD) (38). The input resistance of the PYR soma and dendritic sections 1–5 was 79.84, 80.28, 84.94, 99.36, 137.58, and 246.18 MΩ, respectively (Fig. S2B) based on experimental data (38). The baseline firing rates of the soma and dendritic compartments dend1–dend5 were 2.22 ± 0.98, 2.21 ± 1.27, 2.32 ± 1.2, 2.19 ± 1.16, 2.2 ± 1.4, and 2.67 ± 1.59 Hz (mean ± SD), respectively (Fig. S2D). The F–I curve (firing frequency vs. injected current) of PYRs (Fig. S2E) closely matched previous experimental results (39). Baseline PYR noise was modelled using NEURON’s “INGAUSS” mechanism with mean ± SD = 0 ± 0.092, which generates action potentials at a specified frequency. For a concise overview of these parameters, refer to Table 1.

**Table 1.**
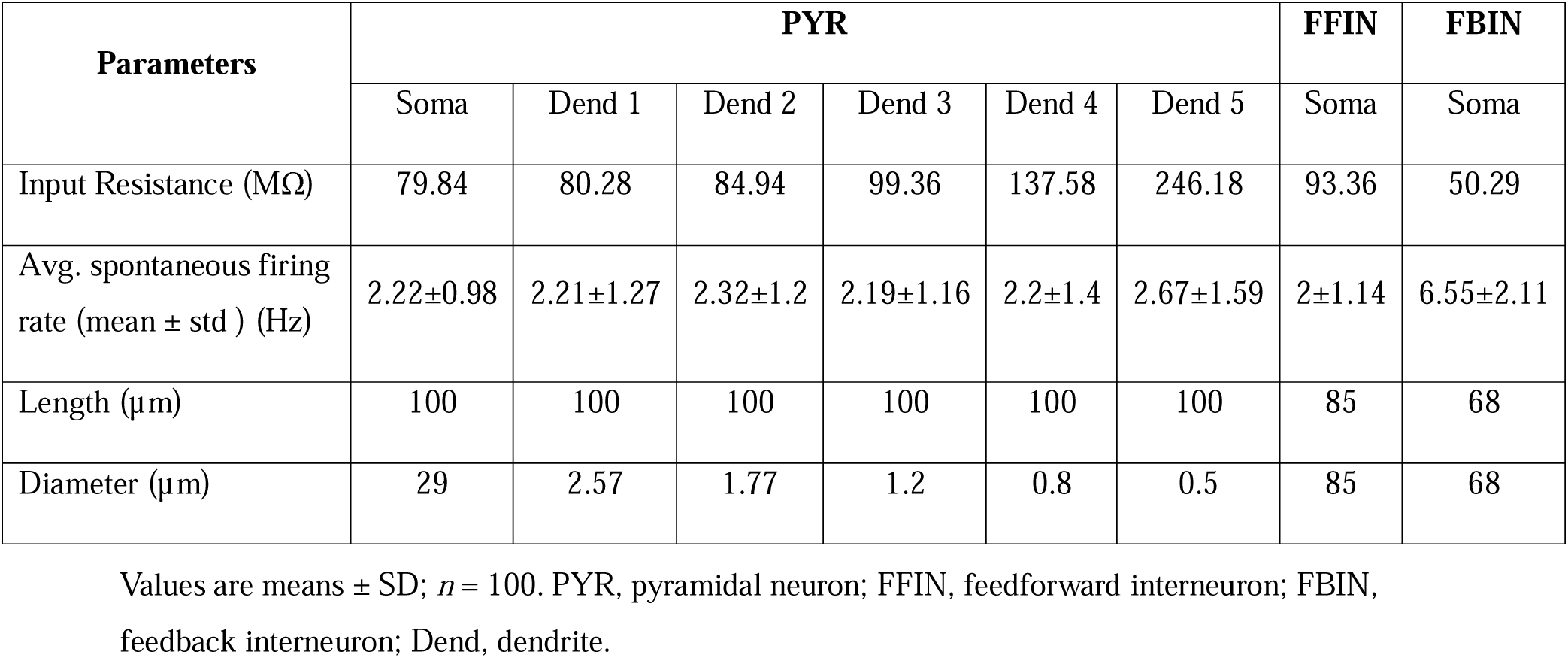
Morphological and biophysical parameters of the modeled cells of the piriform cortex.

#### Feedforward (FF) and Feedback (FB) interneuron model

FFINs and FBINs were modelled as single somatic compartments. The length and diameter of FFINs were both 85 µm, while those of FBINs were 68 µm (Fig. S1B–C). The membrane dynamics of both neuronal types were governed by Equation (1). FFIN and FBIN ion channel mechanisms were adapted from Wang and Buzsáki (1996). For FFINs, the maximum conductance of sodium, potassium, and leak channels was 0.05 S/cm^2^, 0.015 S/cm^2^, and 0.000048 S/cm^2^, respectively. The reversal potentials for sodium, potassium, and leak channels were 55 mV, −90 mV, and −72 mV, respectively. For FBINs, the maximum conductance of sodium, potassium, and leak channels was 0.05 S/cm^2^, 0.01 S/cm^2^, and 0.00014 S/cm^2^, respectively. The reversal potentials for sodium, potassium, and leak channels were 55 mV, −90 mV, and −78 mV, respectively. The resting membrane potentials of FFINs and FBINs were −72 ± 5.9 mV and −77.8 ± 11.3 mV (mean ± SD), respectively (41). The estimated input resistances of FFINs and FBINs were 93.36 MΩ (Fig. S3B(III)) and 50.29 MΩ (Fig. S4B(III)), respectively (41). The current–firing frequency (F–I) curves obtained for FFINs (Fig. S3D) and FBINs (Fig. S4D) corroborated previous experimental results (41). Baseline noise in FFINs and FBINs was modelled using NEURON’s “INGAUSS” mechanism with 0 ± 2.3 and 0 ± 5 (mean ± SD), respectively. The baseline firing rates of FFINs and FBINs were 2.00 ± 1.14 Hz (Fig. S3F) and 6.55 ± 2.11 Hz (Fig. S4F), respectively. Simulations were conducted at a temperature of 33°C.

The somatic compartment of each cell type acts as an integration site for all inputs and generates spikes. This NEURON implementation was based on spike-triggered synaptic transmission in which a spike detected at the presynaptic terminal by a NetCon object causes conductance change at the postsynaptic terminal. A NetCon object connects a presynaptic variable (e.g., membrane voltage) to a target synapse with specified delay and weight parameters (32). Every synapse was modelled using a two-state kinetic scheme defined by the rise time constant Tau1, the decay time constant Tau2, and the reversal potential E_N_. All synapses were modelled using NEURON’s in-built “Exp2Syn” function. Synaptic latencies across different connection types were consistent with findings from Suzuki and Bekkers (2011). The synaptic parameters (Tau1 and Tau2) and weights were adjusted to provide the best fit for target EPSP and IPSP, as per existing literature (36,43). Refer to Table 2 for the comprehensive list of parameters.

**Table 2.** Model parameters PYR, pyramidal neuron; Dend, dendrite; FFIN, feedforward interneuron; FBIN, feedback interneuron; M/T, mitral and tufted cells.

| Synaptic Connections | Synaptic Weight | EPSP/IPSP (mV) | Tau1 (ms) | Tau2 (ms) | $E_N$ (mV) | Synaptic Latency (ms) | Connection Probability |
| --- | --- | --- | --- | --- | --- | --- | --- |
| PYR–PYR Dend 1 | 0.00002 | 0.12 | 0.24 | 3.6 | 0 | 1.54 | 0.2 |
| PYR–PYR Dend 2 | 0.00002 | 0.11 | 0.24 | 3.6 | 0 | 1.54 |  |
| PYR–PYR Dend 3 | 0.00002 | 0.11 | 0.24 | 3.6 | 0 | 1.54 |  |
| PYR–PYR Dend 4 | 0.00002 | 0.11 | 0.24 | 3.6 | 0 | 1.54 |  |
| FBIN–PYR Soma | 0.007 | 1.9 | 0.2 | 1.2 | –80 | 0.65 | 0.35 |
| FFIN–PYR Dend 4 | 0.007 | 1.4 | 0.2 | 1.2 | –80 | 0.6 | 0.3 |
| FFIN–PYR Dend 5 | 0.007 | 0.95 | 0.2 | 1.2 | –80 | 0.6 |  |
| FBIN–FBIN | 0.06 | 1.5 | 0.2 | 1.2 | –80 | 1.54 | 0.1 |
| PYR–FBIN | 0.0001 | 0.31 | 0.26 | 9 | 0 | 0.65 | 0.18 |
| M/T–PYR Dend 5 | 0.000037 | 0.19 | 0.2 | 1.2 | 0 | 1.27 | 0.2 |
| M/T–FFIN | 0.00015 | 0.59 | 0.2 | 1.2 | 0 | 2.42 | 0.4 |
| FFIN–FFIN | 0.0095 | 3 | 0.2 | 1.2 | –80 | 1.54 | 0.4 |

### Odorant Stimulation

To stimulate the PCx network model, we used previously recorded *in vivo* spiking activities from individual mitral and tufted cells (M/Ts) as described in Dasgupta et al., 2022 (18). The OB activity corresponded to three different odor mixtures (stimulus 1: ethyl butyrate and 2-hexanone (1:1); stimulus 2: isopentyl acetate and eucalyptol (1:1); and stimulus Mix: stimuli 1 and 2 (1:1)) presented at two different frequencies (2 Hz and 20 Hz). The 2 Hz frequency lies within the dominant basal sniffing range, whereas 20 Hz represents the higher limit of physiological sniff frequency in awake behaving mice (44,45). Each odorant condition was simulated for 6 s: 2 s baseline, 2 s odor stimulation, and 2 s post-stimulation. For each odor condition, 336 trials were simulated, while synaptic connectivity within the PCx network remained unchanged. Synaptic inputs from M/Ts to PYRs and FFINs were randomized across trials to introduce trial-to-trial variability. Baseline spiking activity across piriform neurons introduced additional trial-to-trial variability and stochasticity in the model.

### Statistical Analysis

Statistical analyses were performed using MATLAB and R. To assess the network-level responses, spiking activity from all 500 pyramidal neurons (PYRs), the principal excitatory output neurons of the piriform cortex, was analyzed across 336 trials for each odor condition (stimulus 1, stimulus 2, and stimulus Mix) presented at 2 Hz and 20 Hz. Spike times were binned using a 50 ms sliding time window. For each trial, average firing rates were computed for individual neurons within two post-stimulus time windows (0–0.5 s and 0–2 s). The early 0–0.5 s window, corresponding to one cycle of the slower 2Hz stimulus frequency, was selected to avoid confounding effects arising from differences in the total odor delivery across frequencies, while also matching the decision-making time based on previously observed behavioral assays (18). The baseline firing rate for each neuron was calculated within the pre-stimulus window (−1.5 to −0.5 s) and subtracted from the stimulus-evoked firing rate on a per-trial basis, hereafter referred to as baseline correction.

#### Violin plots

Two sets of comparisons were performed: (1) odor identity comparisons (e.g., stimulus 1 vs. Mix, stimulus 2 vs. Mix, and stimulus 1 vs. stimulus 2) for each frequency (2 Hz and 20 Hz); and (2) odor frequency comparisons (2 Hz vs. 20 Hz) for each odor type (1, 2, and Mix). For each odor condition (stimuli 1, 2, and Mix at 2 Hz and 20 Hz), baseline-corrected firing rates were averaged across 336 trials per neuron to generate firing rate distributions across all PYRs. Odor identity comparisons were then performed pairwise within the 0–0.5 s stimulus window, while odor frequency comparisons were performed pairwise within two stimulus time windows (0–0.5 s and 0–2 s).

Prior to statistical testing, data distributions were assessed for normality using the Lilliefors test. As the firing rate distributions across all 500 PYRs significantly deviated from normality (*p* < 0.05), non-parametric Mann–Whitney U tests were performed for both comparison types. Cliff’s delta (δ) (46,47) was used to quantify the non-parametric effect size between distributions. This was calculated as the difference between number of instances (#) where values from Group 1 (x_i1_) are larger than Group 2 (x_j2_), and number of instances (#) where values from Group 1 (x_i1_) are smaller than Group 2 (x_j2_), divided by the product of sample sizes of Group 1 (n_1_), and Group 2 (n_2_), as follows:

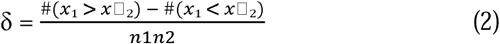

Effect size magnitudes were interpreted using the following thresholds: |δ| < 0.147 as negligible, |δ| < 0.33 as small, |δ| < 0.474 as medium, and |δ| ≥ 0.474 as large (48). Cliff’s δ was estimated using the ‘effsize’ package in R. To evaluate whether odor-evoked changes in the population firing rates differed across genotypes (Control vs. Knockout) and stimulation frequencies (2 Hz vs. 20 Hz), a two-way mixed-design repeated-measures ANOVA with the Aligned Rank Transform (ART) (49,50) procedure was implemented using the ‘*ARTool*’ package in R. ART is a non-parametric procedure used to assess both main and interaction effects in factorial designs. ART preprocesses the data by first aligning it (51,52) and then applying mid-ranks (53) to the aligned data. The resulting aligned-and-ranked data were then analyzed using ANOVA. For these analyses, genotype was treated as a between-subject factor, whereas stimulation frequency as a within-subject factor, with neuron ID included as a random effect.

#### Analysis of single neuronal parameters

Input resistance was calculated from the slope of the steady-state voltage–current relationship obtained using subthreshold current steps. Action potential threshold was defined as the membrane potential (V_m_) at which dV_m_/dt first exceeded 20 V/s. Baseline firing rate for each neuron type was calculated as the mean ± SD firing rate from single-cell simulations driven by synaptic noise over a duration of 1.5 s across 100 trials. F–I plots were generated by quantifying firing frequency as a function of injected current over 1 s without noise being added. EPSP and IPSP for each type of synaptic connection were determined as the peak depolarisation or hyperpolarisation from the resting membrane potential (RMP), respectively.

#### One dimensional convolutional neural network (1D-CNN) model

We employed 1D convolutional neural network (CNNs) to perform binary classification of odor stimuli based on PYRs’ spiking activity (Fig. 2A). For this, spike trains were binned into non-overlapping 20 ms windows, producing binary activity vectors indicating whether a neuron spiked at least once within the specified time bin. Thus, the dataset dimensions for any binary classification task were *672 × 100 × N* where 672 *(2 × 336)* was the number of trials, *N* was the size of the PYR population used for the particular classification, and 100 was the number of time bins (2000/20). Each dataset was partitioned into training and testing sets, with 60% and 40% of the data, respectively. The input to the 1D-CNN model consisted of matrices of size *100 × N*. In the 1D-CNN architecture, convolution was applied along the temporal dimension. The convolutional layer applied temporal filters (kernel size = 3) across the input to extract time-dependent features. Convolution is a linear operation that involves multiplying the inputs with a set of weights. In our case, inputs were multiplied with the two-dimensional array of weights, known as a filter, of dimension *3 × N* where 3 is the kernel size. This operation assigns a unique value to each pass, resulting in a feature map of dimension 100. Once computed, each value in the feature map was passed to the rectified linear unit (ReLU) activation function. ReLU activation function transforms negative values to zero while retaining positive values. Convolutional neural networks are susceptible to overfitting, so a regularization technique that sets the inputs to 0 for some fraction of neurons during the training phase was used. Finally, the outputs from the 1D convolutional layer were flattened into a one-dimensional vector using a flatten layer before the fully connected dense layers. These dense/fully connected layers, after application of the activation function (Softmax), map the outputs to the class probability distributions. Our 1D-CNN models consisted of 2 convolution layers and 2 dense layers with a dropout rate of 0.3 applied after each layer as shown in Fig. 2A. The first convolutional layer consisted of 128 filters with a kernel size of 3 along with the ReLU activation function. This layer transformed the input matrix of size *100 × N* into feature maps of size *100 × 128*. A dropout rate of 0.3 was applied after each layer to mitigate overfitting. It was followed by a second convolutional layer consisting of 128 filters with a kernel size of 3. The output from this layer was then flattened into a one-dimensional vector. Finally, two dense layers with ReLU activation were followed by a final dense layer with the Softmax activation function for predicting the probability of the target stimulus. The model was trained using the Adam optimizer for weight optimization and categorical cross-entropy as the loss function. The model was trained for 75 epochs with a batch size of 32. To prevent overfitting and unnecessary training, early stopping was implemented based on validation loss, with a patience parameter of 5 epochs. We conducted a series of experiments to identify the parameters that optimized the CNN’s classification performance.

**Figure 1.**
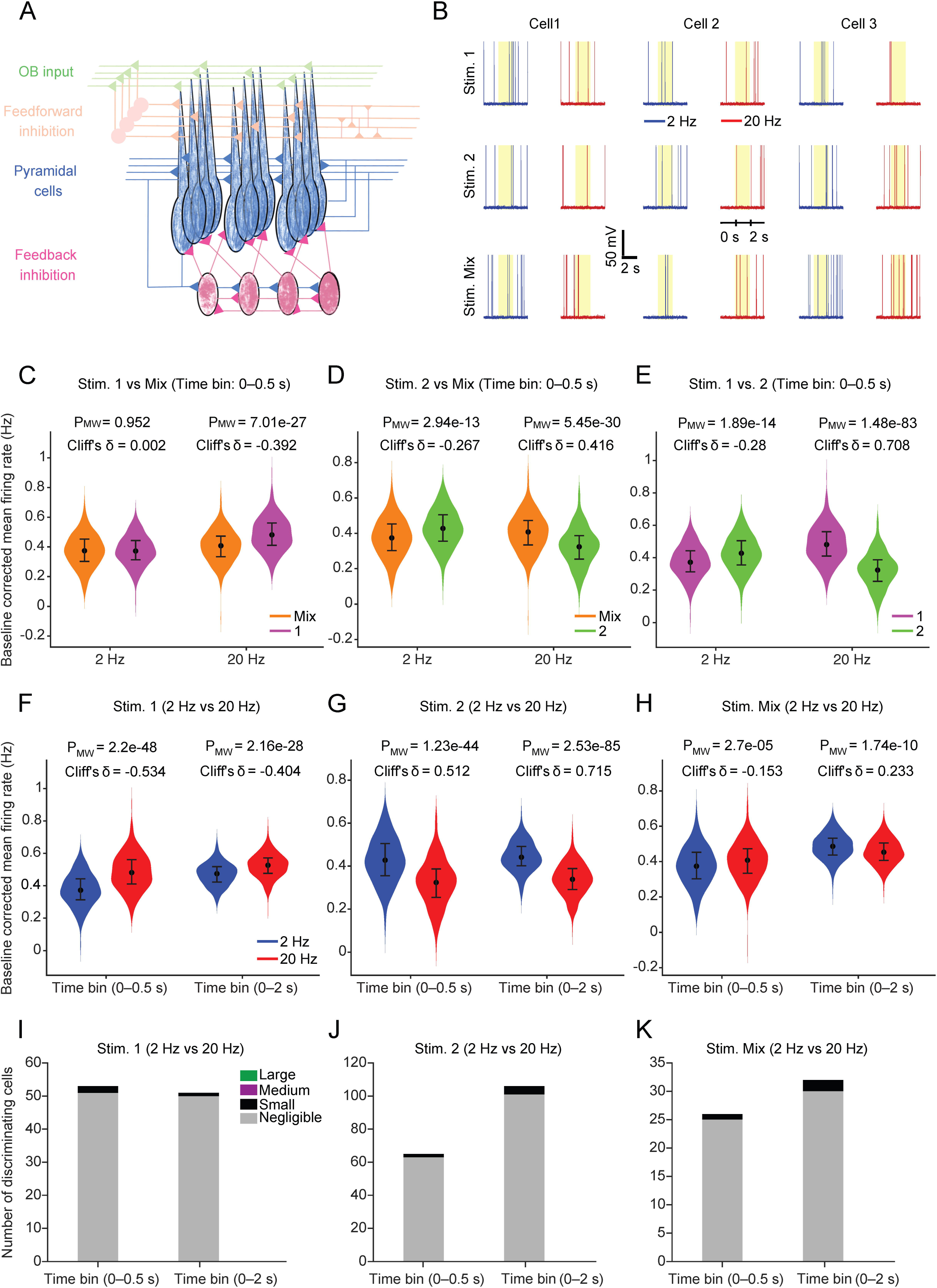
PCx neurons discriminate between odor identities and frequencies at the network level. **A,** Schematic of the piriform cortex (PCx) network model. **B,** Representative voltage traces from three model pyramidal neurons (PYRs) in response to the three stimuli (1, 2, and Mix) presented at 2 Hz (blue) and 20 Hz (red) frequencies. **C,** Violin plots showing the distribution of mean firing rates (averaged across 336 trials per neuron) after baseline subtraction during the initial 500 ms of odor stimulation at 2 Hz and 20 Hz: stimulus 1 vs. Mix (**D**), stimulus 2 vs. Mix, and (**E**) stimulus 1 vs. stimulus 2; statistical significance was tested using two-tailed Mann–Whitney U test, and effect sizes were calculated using Cliff’s delta (δ) (*n* = 500 neurons). In each violin plot, the central solid marker represents the median while the whiskers represent the first and third quartiles. **F,** Violin plots showing the distribution of mean firing rates (averaged across 336 trials per neuron) after baseline subtraction in response to stimulus 1 at 2 Hz vs. 20 Hz, **(G),** stimulus 2 at 2 Hz vs. 20 Hz, and **(H),** stimulus Mix at 2 Hz vs. 20 Hz. The time bins represent the odor stimulation period used for the analysis (0–0.5 s and 0–2 s). Statistical tests were performed in a similar way as described in Fig. 1C–E. **I,** Stacked bar plots depicting effect size magnitudes among pyramidal neurons with significant firing rate differences between 2 Hz and 20 Hz stimulation (*p* < 0.05, two-tailed Mann–Whitney U test comparing 336 trials per frequency per neuron) for stimulus 1, **(J),** stimulus 2, and **(K),** stimulus Mix during the 0–0.5 s and 0–2 s post-stimulus windows. Each bar represents discriminating neurons stratified by Cliff’s delta (δ) magnitude: negligible (|δ| < 0.147; grey), small (0.147 ≤ |δ| < 0.33; black), medium (0.33 ≤ |δ| < 0.474; purple), and large (|δ| ≥ 0.474; green). Note that none of the conditions showed any cells with medium or large magnitude differences.

**Figure 2.**
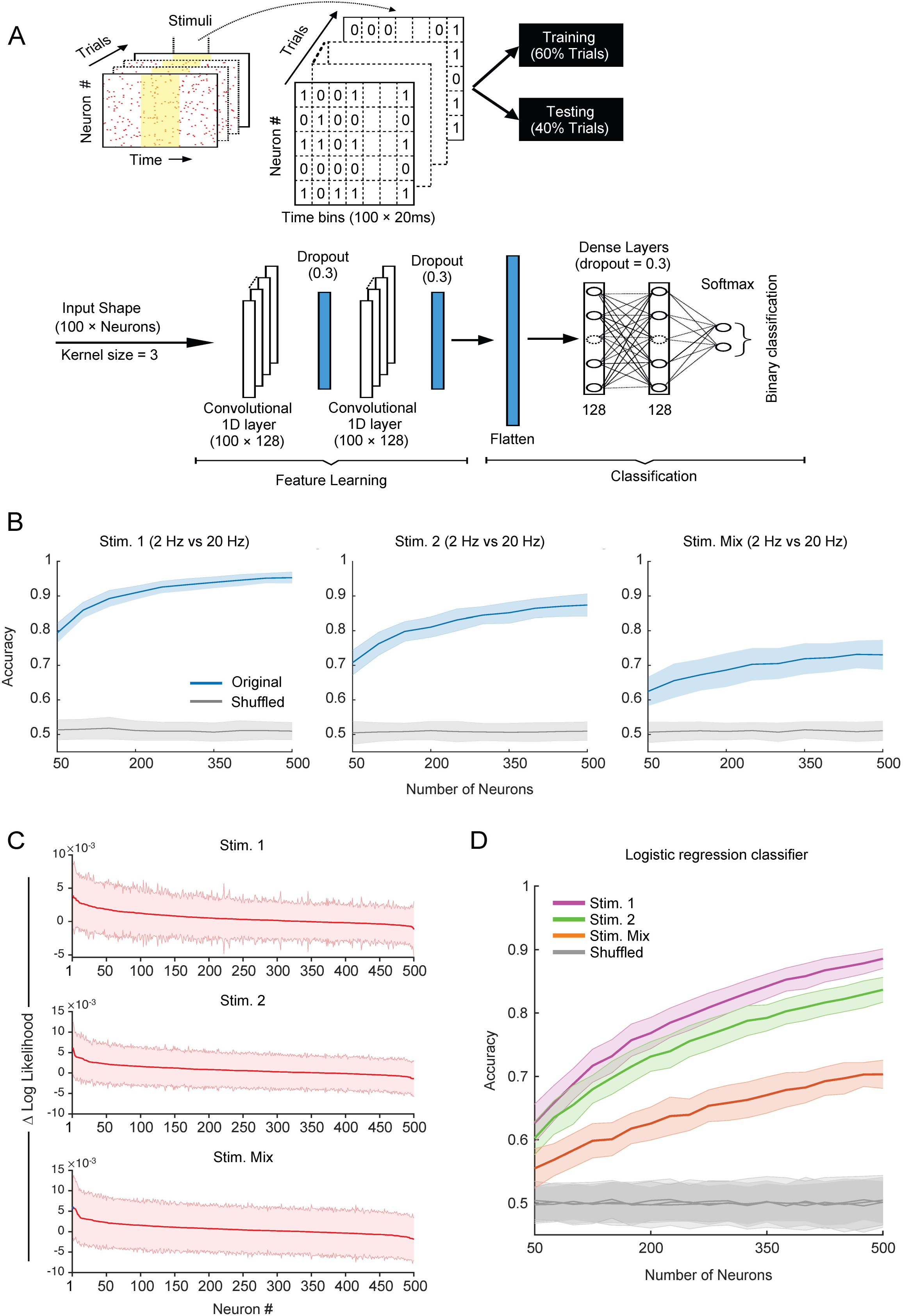
One-dimensional convolutional neural network models (1D-CNNs) and logistic regression classifiers demonstrate that PCx network activity can discriminate odor frequencies. **A,** Schematic showing the preparation of datasets using spike times of PYRs during the entire 2 s odor stimulation period and the configuration of 1D-CNN for odor-frequency classification. **B,** Average classification accuracy of CNN models in discriminating 2 Hz vs. 20 Hz stimulation for stimulus 1 (left), stimulus 2 (center), and stimulus Mix (right) as a function of PYR population size (population step size = 50). The solid line and shaded area represent the mean and the corresponding standard deviation, respectively, from 144 iterations of randomly sampled neurons. Chance-level accuracy was estimated by randomly shuffling trial labels (grey lines). **C,** Plots showing the difference in log-likelihood values (ΔLL) of CNN model between intact and perturbed test sets for each neuron in the descending order of their importance for frequency decoding (2 Hz vs. 20 Hz) for stimulus 1 (top), stimulus 2 (middle), and stimulus Mix (bottom). In the perturbed test set, a neuron’s spiking activity was replaced by its baseline activity. The solid lines and the shaded region have similar meaning as in Fig. 2B (*n* = 300 iterations). **D,** Average classification accuracy of logistic regression classifiers in discriminating 2 Hz vs. 20 Hz stimulation for stimulus 1 (magenta), stimulus 2 (green), and stimulus Mix (orange) as a function of PYR population size (population step size = 25). For each population size, neurons were randomly sampled in 144 independent iterations. Solid lines represent the mean, and the shaded region represents the corresponding standard deviation. Chance-level accuracy was estimated by randomly shuffling trial labels (grey lines).

#### Logistic regression classifier

The data from all network models were also trained and tested using Scikit-learn’s SGD classifier with a logistic loss function to decode odor stimulation frequencies (2 Hz vs. 20 Hz) for each stimulus (1, 2, and Mix). As with the CNN models, these classifiers were also used to evaluate how odor frequency decoding accuracy scaled with the population size of pyramidal neurons (PYRs). Data preparation for binary classification of each odor type was identical to that used for the CNN models. For each trial, spike trains of all PYRs recorded during the 2 s odor stimulation period were divided into 100 non-overlapping time bins of 20 ms each. Each time bin was assigned a value of 1 if at least one spike occurred within that bin for a given neuron, and 0 otherwise. This procedure resulted in a binary matrix representation of neural activity for a single trial with dimensions *100 × N*, where *N* denotes the number of neurons. Across trials, these matrices were stacked to form a dataset of size *672 × 100 × N* (336 trials per frequency condition).

For training and testing, the dataset was split into 60% and 40% of trials, respectively. Each trial’s activity matrix was flattened into a one-dimensional feature vector of length *100 × N*, which served as input to the classifier. Prior to training, all feature vectors were standardized using Scikit-learn’s StandardScaler, ensuring zero mean and unit variance. The classifier was implemented using Scikit-learn’s Pipeline framework, consisting of the StandardScaler followed by an SGDClassifier configured with a logistic (log) loss function, corresponding to logistic regression. The stochastic gradient descent (SGD) optimizer updated the model weights iteratively to minimize the logistic loss, offering computational efficiency and scalability for high-dimensional data. The model was trained for up to 1000 iterations with a convergence tolerance of 1 × 10^−3^.

To assess how decoding accuracy scales with the PYR population size, the classifier was trained and tested on randomly sampled subsets of the PYR population. The number of neurons per subset ranged from 50 to 500 neurons, in increments of 25. For each population size, 144 independent iterations were performed using different randomly sampled neuronal subsets, and the mean accuracy ± standard deviation across iterations was calculated. To establish a baseline for chance-level performance, a shuffle control was conducted in parallel for each iteration. In this control, the class labels in the training data were randomly shuffled while keeping the feature matrices unchanged, thereby disrupting the true relationship between neural activity and stimulus frequency. The resulting shuffled accuracy represented chance-level decoding performance, against which classifier performance was compared.

#### Accuracy vs. cell number

To quantify how decoding performance scales with neuronal population size, 1D-CNNs were trained and tested on PYRs subsets ranging from 50 to 500 neurons in increments of 50. For statistical robustness, 144 independent trials were conducted per population size. In each iteration, neurons were randomly sampled from the entire PYR population, and data (i.e., *2 × 336* trials) were randomly split into training (60%) and testing (40%) sets. To establish a baseline for chance-level performance, we trained the model on datasets where the stimulus labels were randomly shuffled. This disrupted any real relationship between spiking patterns and stimulus frequency while preserving the overall distribution of spike data. Decoding accuracy on the test set was recorded for each trial/iteration under both actual and shuffled-label conditions. Finally, decoding accuracies were averaged across trials to compute mean accuracy and standard deviation for each population size. This analysis quantified the marginal improvement in decoding accuracy with increasing population size and determined whether decoding performance plateaued at larger population sizes.

#### Determining cell importance

The importance of individual neurons for binary classification was quantified using negative cross-entropy loss (log-likelihood) of the CNN model. Log-likelihood measures how well the model’s probability predictions align with the true class labels (1 or 0). Mathematically, it is calculated as the sum of log predicted probabilities of true class labels in the test set. Higher values of log-likelihood indicate better model performance, meaning stronger alignment between predictions and actual class labels. PYRs odor frequency trials were randomly divided into training (70%) and test (30%) sets. The CNN models were trained and tested on the data corresponding to the odor stimulation period (i.e., 2–4 s) and the log-likelihood was calculated. We then replaced the stimulus-evoked activity of a given neuron with its baseline/pre-stimulus activity in the test set and again calculated the log-likelihood. The impact of this perturbation was quantified as the difference in the model log-likelihood between intact and perturbed conditions (ΔLL). A substantial reduction in ΔLL indicated that the neuron contributed strongly to decoding performance. We performed this analysis for all 500 PYRs one at a time. The neuronal importance score was defined as the mean ΔLL across 300 iterations, with higher values indicating greater contribution to stimulus discrimination. Finally, neurons were ranked by their mean importance scores, thereby revealing their relative contribution to the model performance. This procedure was performed separately for each experimental condition (stimuli 1, 2, and Mix). CNN architecture for this analysis was largely the same as outlined above, except that the number of training epochs was reduced to 15 to minimize overfitting.

#### Accuracy using baseline firing data

To evaluate model behavior in the absence of odor stimulation, we trained and tested the model exclusively on baseline firing activity (0–2 s) of all 500 PYRs using the same CNN architecture and method outlined above. This analysis tested whether the model learned any spurious patterns not directly related to the stimulus (Fig. S5). A total of 144 iterations were conducted, with each iteration using a random 60%/40% train–test split.

#### PYR population baseline firing rate

Spike data from all 500 pyramidal neurons were binned into 2 s time windows and converted to firing rates (Hz) for all six odor conditions across 336 trials. Population baseline firing rates across all network conditions were calculated by averaging firing rates during the first time bin (0–2 s, i.e., baseline period) across all neurons, trials, and odor conditions (Fig. 9B). Data were analyzed using custom MATLAB functions, with values reported as mean ± standard deviation across all network conditions.

## Results

To investigate whether odor frequency is encoded in the piriform cortex (PCx), we simulated a biophysically relevant network model of PCx comprising 500 pyramidal neurons (PYRs), 50 feedforward interneurons (FFINs), and 50 feedback interneurons (FBINs) (Fig. 1A). The model architecture and connection probabilities were based on experimental data from rodents (43,54,55), as described in the Materials and Methods section. Previously recorded *in* vivo M/T cells activities from Dasgupta et al., 2022 (18) served as the olfactory bulb (OB) output and was used as input to the PCx model. OB activity was recorded in response to three odor stimuli mixtures (stimulus 1: ethyl butyrate and 2-hexanone (1:1); stimulus 2: isopentyl acetate and eucalyptol (1:1); stimulus Mix: stimuli 1 and 2 (1:1)) each presented at two frequencies (2 Hz and 20 Hz). The model was simulated using the NEURON environment for 6 s (2 s baseline followed by 2 s of odor stimulation and 2 s post-stimulation).

Each PYR was modelled with a soma connected to an apical dendritic trunk consisting of five dendritic sections (Fig. S1A). The FFINs and FBINs were each modelled as single somatic compartment (Fig. S1B–C). Resting membrane potentials of PYRs, FFINs, and FBINs were −74.1 ± 8.6 mV, −72 ± 5.9 mV, and −77.8 ± 11.3 mV, respectively, consistent with previous experimental findings (38,41). Action potential firing profiles of PYRs, FFINs, and FBINs (Fig. S2E; Fig. S3D; Fig. S4D) closely matched experimentally observed recordings (39,41). Spontaneous baseline firing rates at the soma of PYR, FFIN, and FBIN were 2.22 ± 0.98 Hz (Fig. S2D), 2 ± 1.14 Hz (Fig. S3F), and 6.55 ± 2.11 Hz (Fig. S4F), respectively (56). The estimated input resistance values measured from the soma and the dendrites of PYRs (79.84, 80.28, 84.94, 99.36, 137.58, and 246.18 MΩ from the soma and dend1 to dend5, respectively) (Fig. S2B) were consistent with experimental findings (38). The input resistance measured from the FFINs and the FBINs was 93.36 MΩ (Fig. S3B(III)) and 50.29 MΩ (Fig. S4B(III)), respectively (41). Spike timings of all 500 PYRs were recorded across 336 trials for each odor stimulation condition. Spike trains from individual trials were converted to firing rates within two post-stimulus windows: the initial 0–0.5 s and the entire 0–2 s of the odor stimulation period.

### Pyramidal neurons of the PCx can largely discriminate odor identity irrespective of the odor frequency

Overall, we had six odor stimulation conditions: stimulus 1, stimulus 2, and stimulus Mix each presented at 2 Hz and 20 Hz. To investigate whether the PYR population activity could discriminate odor identity, firing rate distributions corresponding to the initial 0–0.5 s window were compared across different odor pairs for the 2 Hz and 20 Hz conditions (Fig. 1C–E). At 2 Hz, the PYR population response to stimulus Mix and stimulus 1 showed a non-significant difference (Mix: 0.37 Hz, IQR = 0.30–0.45 Hz, *n* = 500, and stimulus 1: 0.37 Hz, IQR = 0.31–0.44 Hz, *n* = 500; Mann–Whitney *p* = 0.952; Cliff’s |δ| = 0.002, “negligible” effect size; Fig. 1C). At 20 Hz, however, we observed a significant difference between these distributions (Mix: 0.41 Hz, IQR = 0.33–0.47 Hz; stimulus 1: 0.48 Hz, IQR = 0.41–0.56 Hz; Mann–Whitney *p* = 7.01e−27; Cliff’s |δ| = 0.392, “medium” effect size; Fig. 1C). For stimulus Mix vs. stimulus 2, PYRs exhibited small but significant differences in population-level firing at 2 Hz (Mix: 0.37 Hz, IQR = 0.30–0.45 Hz, *n* = 500; stimulus 2: 0.43 Hz, IQR = 0.36–0.50 Hz, *n* = 500; Mann–Whitney *p* = 2.94e−13; Cliff’s |δ| = 0.267, “small” effect size; Fig. 1D), while at 20 Hz, PYR population firing rates were significantly higher for stimulus Mix than for stimulus 2 (Mix: 0.41 Hz, IQR = 0.33–0.47 Hz; stimulus 2: 0.32 Hz, IQR = 0.25–0.39 Hz, *n* = 500; Mann–Whitney *p* = 5.45e−30; Cliff’s δ = 0.416, “medium” effect size; Fig. 1D). Comparing stimulus 1 and stimulus 2, responses differed modestly but significantly at 2 Hz (stimulus 1: 0.37 Hz, IQR = 0.31–0.44 Hz; stimulus 2: 0.43 Hz, IQR = 0.36–0.50 Hz, *n* = 500; Mann–Whitney *p* = 1.89e−14; Cliff’s |δ| = 0.28, “small” effect size; Fig. 1E), whereas at 20 Hz, PYRs fired more strongly for stimulus 1 than for stimulus 2 (stimulus 1: 0.48 Hz, IQR = 0.41–0.56 Hz; stimulus 2: 0.32 Hz, IQR = 0.25–0.39 Hz, *n* = 500; Mann–Whitney *p* = 1.48e−83; Cliff’s |δ| = 0.708, “large” effect size; Fig. 1E).

Taken together, these results demonstrate that the PCx model can discriminate between the identities of odor stimulus mixtures at both 2 Hz and 20 Hz stimulation frequencies. Nevertheless, odor identity discrimination was weaker (“*negligible*” to “*small*” effect sizes) at 2 Hz compared to the 20 Hz (“*medium*” to “*large*” effect sizes).

### The piriform cortex network model can discriminate 2 Hz odor frequency from 20 Hz

Next, we examined whether the PCx responds differently to the 2 Hz and 20 Hz stimulation frequencies of the same odor stimulus. PYRs’ responses were quantified over two post-stimulus time windows: the initial 0–0.5 s and the entire 0–2 s odor stimulation period across 336 trials. For each odor, baseline-corrected mean firing rate distributions across the entire PYR population were compared between 2 Hz and 20 Hz stimulation for each time window (Fig. 1F–H).

For stimulus 1, during the 0–0.5 s window, the PYR population exhibited a significant increase in median firing rate from 0.37 Hz (IQR = 0.31–0.44 Hz) at 2 Hz stimulation to 0.48 Hz (IQR = 0.41–0.56 Hz) at 20 Hz stimulation (*n* = 500; Mann–Whitney *p* = 2.2e−48; Cliff’s δ = −0.534, “large” effect size; Fig. 1F). During the 0–2 s window, median firing rate increased from 0.47 Hz (IQR = 0.42–0.52 Hz) at 2 Hz to 0.53 Hz (IQR = 0.48–0.57 Hz) at 20 Hz (*n* = 500; Mann–Whitney *p* = 2.16e−28; Cliff’s δ = −0.404, “medium” effect size; Fig. 1F). For stimulus 2, during the 0–0.5 s window, median firing rate decreased significantly from 0.43 Hz (IQR = 0.36–0.5 Hz) at 2 Hz to 0.32 Hz (IQR = 0.25–0.39 Hz) at 20 Hz (*n* = 500; Mann–Whitney *p* = 1.23e−44; Cliff’s δ = 0.512, “large” effect size; Fig. 1G). Similarly, during the 0–2 s window, the median firing rate also decreased significantly from 0.44 Hz (IQR = 0.40–0.49 Hz) at 2 Hz to 0.34 Hz (IQR = 0.29–0.39 Hz) at 20 Hz (*n* = 500; Mann–Whitney *p* = 2.53e−85; Cliff’s δ = 0.715, “large” effect size; Fig. 1G). For stimulus Mix, PYRs showed a small but significant increase in firing rate during the 0–0.5 s window, from a median of 0.37 Hz (IQR = 0.30–0.45 Hz) at 2 Hz to 0.41 Hz (IQR = 0.33–0.47 Hz) at 20 Hz (*n* = 500; Mann–Whitney *p* = 2.7e−05; Cliff’s δ = −0.153, “small” effect size; Fig. 1H). In contrast, during the 0–2 s window, the median firing rate decreased slightly from 0.49 Hz (IQR = 0.44–0.53 Hz) at 2 Hz to 0.45 Hz (IQR = 0.41–0.51 Hz) at 20 Hz (*n* = 500; Mann–Whitney *p* = 1.74e−10; Cliff’s δ = 0.233, “small” effect size; Fig. 1H).

These findings indicate that the PYR population can discriminate between 2 Hz and 20 Hz odor stimulation frequencies regardless of the odor identity.

However, to determine whether individual neurons can also discriminate odor frequencies, we analyzed the activity of all 500 PYRs individually using the same time windows and trials as before, applying a two-tailed Mann–Whitney U test (*n* = 336). Neurons exhibiting significant differences (*p* < 0.05) were classified as odor frequency-discriminating cells, and effect sizes were estimated using Cliff’s |δ|. For stimulus 1, during the 0–0.5 s window, 53 out of 500 neurons were frequency-discriminating, with 51 and 2 of them showing “negligible” and “small” effect sizes, respectively (Fig. 1I). During the 0–2 s window, 51 out of 500 neurons were frequency-discriminating, with 50 and 1 showing “negligible” and “small” effect size, respectively (Fig. 1I). For stimulus 2, during the 0–0.5 s window, 65 of 500 neurons were frequency-discriminating, of which 63 and 2 cells showed “negligible” and “small” effect sizes, respectively (Fig. 1J). During the 0–2 s window, 106 neurons were frequency-discriminating, with 101 and 5 showing “negligible” and “small” effect sizes, respectively (Fig. 1J). For stimulus Mix, 26 out of 500 neurons were frequency-discriminating during the 0–0.5 s window, with 25 and 1 showed “negligible” and “small” effect sizes, respectively, whereas during the 0–2 s window, 32 neurons were frequency-discriminating, with 30 and 2 showed “negligible” and “small” effect sizes, respectively (Fig. 1K).

Given the multiple comparisons involved in testing 500 individual neurons, a Bonferroni correction was applied (corrected threshold: *p* < 0.05/500). Under this conservative criterion, no frequency-discriminating cells were identified for stimulus 1 and stimulus 2 in either time window. For stimulus Mix, only a single neuron showed significant discrimination during the 0–2 s window (“small” effect size), while no discriminating neurons were found in the 0–0.5 s window.

Collectively, these findings suggest that spike count responses of individual PYRs are insufficient to reliably distinguish odor frequencies. In contrast, the PYR population as a whole showed significant frequency discrimination, suggesting that this capacity likely emerges at the population level through coordinated activity across neurons.

To examine how accurately odor frequency could be decoded from the PYR population activity, PYR responses were quantified as binary spike vectors (1 s and 0 s) over the entire 2 s odor stimulation period. Spiking activity from each trial was divided into 20 ms non-overlapping time bins, where a value of 1 indicated at least one spike within that bin. Sixty percent of the total activity matrices from 2 Hz and 20 Hz odor trials were used to train one-dimensional convolutional neural network (1D-CNN) models, which were then tested on the remaining 40% of the trials (Fig. 2A).

The 1D-CNN models achieved peak decoding accuracies of 0.95 ± 0.016, 0.87 ± 0.032, and 0.73 ± 0.043 (mean ± SD) for distinguishing 2 Hz odor stimulation from 20 Hz for stimuli 1, 2, and Mix, respectively. Shuffled controls yielded chance-level peak accuracies of 0.51 ± 0.025, 0.50 ± 0.027, and 0.51 ± 0.028 for stimuli 1, 2, and Mix, respectively. To evaluate how decoding accuracy scaled with population size, CNN models were trained and tested on randomly selected subsets of PYR population of increasing size, each evaluated over 144 independent iterations (Fig. 2B). The classification accuracy increased monotonically with the number of neurons included and consistently outperformed shuffled controls across all odor conditions. To ensure that the CNN models were free of any intrinsic bias, additional models were trained and tested on the baseline activity of PYRs in an identical manner (Fig. S5), which showed no significant deviation from their corresponding shuffle controls.

We next quantified the relative contribution of individual pyramidal neurons to CNN decoding accuracy. To this end, in the test set, each neuron’s odor-period activity was systematically replaced with its baseline activity, one neuron at a time, and the log-likelihood was recalculated. For each perturbation, the mean change in log-likelihood (ΔLL; intact minus perturbed) was computed across 300 independent iterations, and neurons were ranked in descending order of ΔLL (Fig. 2C). Larger positive ΔLL values indicated greater contribution to odor frequency discrimination, while near-zero or negative values reflected minimal or adverse contributions. We observed that the ΔLL values for all three odors are distributed along a continuum. Neurons at the left extreme contribute positively, those at the right extreme contribute negatively, and those in between contribute equally and minimally. Taken together, these findings suggest that the odor frequency discrimination capacity is distributed across the entire PYR population.

To validate the non-linear model (CNN) findings using a linear model, logistic regression classifiers were trained and tested to decode odor stimulation frequency for each odor stimulus type (stimuli 1, 2, and Mix; Fig. 2D). Data preparation for this linear classifier was identical to that used for the CNN models, except that the inputs were formatted as one-dimensional feature vectors of length *100 × N*, where 100 represents the number of 20 ms time bins, and *N* is the number of neurons. The logistic regression classifiers achieved mean peak accuracies of 0.89 ± 0.015, 0.84 ± 0.019, and 0.70 ± 0.022 for stimuli 1, 2, and Mix, respectively. Decoding performance of the logistic regression models increased monotonically with population size (Fig. 2D) and closely resembled the performance of CNN models (Fig. 2B).

Together, these findings indicate that the spiking patterns of the PYR population carry crucial information for odor frequency discrimination. The fact that both a non-linear (1D-CNN) and a linear (logistic regression) decoder successfully captured this information suggests that odor frequency encoding is a network-level phenomenon with a distributed role of the PYRs.

### Different synaptic connections differentially affect odor frequency discrimination

Synaptic inhibition, mediated by local inhibitory microcircuits, plays a crucial role in shaping information processing in the central nervous system (57–59). In the mammalian olfactory bulb, GABAergic interneurons are crucial for regulating the precise spike timing and synchrony of mitral and tufted cells (M/Ts) through lateral and recurrent inhibition (60–63). The PCx receives dense and distributed projections from M/T axon collaterals and contains extensive recurrent excitatory connections among pyramidal neurons. Because of this highly recurrent architecture, synaptic inhibition is essential for maintaining the excitation/inhibition balance required for stabilizing the network activity (64). Furthermore, inhibition in the olfactory bulb of mice and the antennal lobe of insects and honeybees has been reported to modulate odor identity discrimination in a bidirectional way (65–73). Moreover, the local inhibition has been shown to bidirectionally shift the coupling between odor frequency and membrane potential fluctuations of M/Ts in the olfactory bulb (18). On the other hand, inhibition in the PCx has been reported to play a significant role in sparse odor representations (35,74), gain control (75), concentration-invariant odor coding (56), and network oscillations (35,76–78).

However, little is known about how circuit imbalance in the PCx influences olfactory coding of naturalistic odor stimuli, especially the temporal features. Synaptic inhibition in the PCx is mediated by feedforward and feedback interneuron microcircuits, while excitation is largely driven by recurrent connections among pyramidal neurons. Therefore, we investigated how synaptic perturbations in each of these canonical circuit motifs would affect odor frequency discrimination. To this end, we systematically removed each synaptic connection type from the control network model, one at a time, and assessed its effect on odor frequency discrimination. For each knockout condition, the PCx knockout models were simulated and analyzed using the same approach as the control network.

### Eliminating either feedforward or feedback inhibition enhanced odor frequency discrimination

To investigate the influence of direct inhibitory inputs onto PYRs in the context of odor frequency discrimination, we selectively removed the synaptic connections from feedforward (FF), and feedback (FB) interneurons onto PYRs, one at a time (Fig. 3A; Fig. 4A). We then trained 1D-CNN models as described previously to classify 2 Hz vs. 20 Hz odor trials based on PYR population activity. Both knockout models yielded higher decoding accuracy than the control network across all population sizes (Fig. 3B; Fig. 4B; Fig. S6). For the FF–PYR knockout model, CNNs achieved peak decoding accuracies (mean ± SD, *n* = 500 neurons) of 0.98 ± 0.006, 0.96 ± 0.011, and 0.87 ± 0.03 for stimulus 1, 2 and Mix respectively (Fig. 3B). Logistic regression classifiers trained on the same activity achieved peak accuracies of 0.97 ± 0.01, 0.94 ± 0.015, and 0.81 ± 0.019, respectively (Fig. S6). Similarly, for the FB–PYR knockout model, CNNs achieved peak accuracies of 0.99 ± 0.005 for stimulus 1, 0.97 ± 0.011 for stimulus 2, and 0.88 ± 0.02 for stimulus Mix (Fig. 4B). Logistic regression classifiers again produced comparable peak accuracies of 0.98 ± 0.007, 0.97 ± 0.009, and 0.88 ± 0.02, respectively, for the same stimuli (Fig. S6). Beyond machine learning-based decoding, odor frequency discrimination was further assessed using distributions of baseline-corrected mean firing rates across the PYR population and compared with the control using ART-ANOVA.

**Figure 3.**
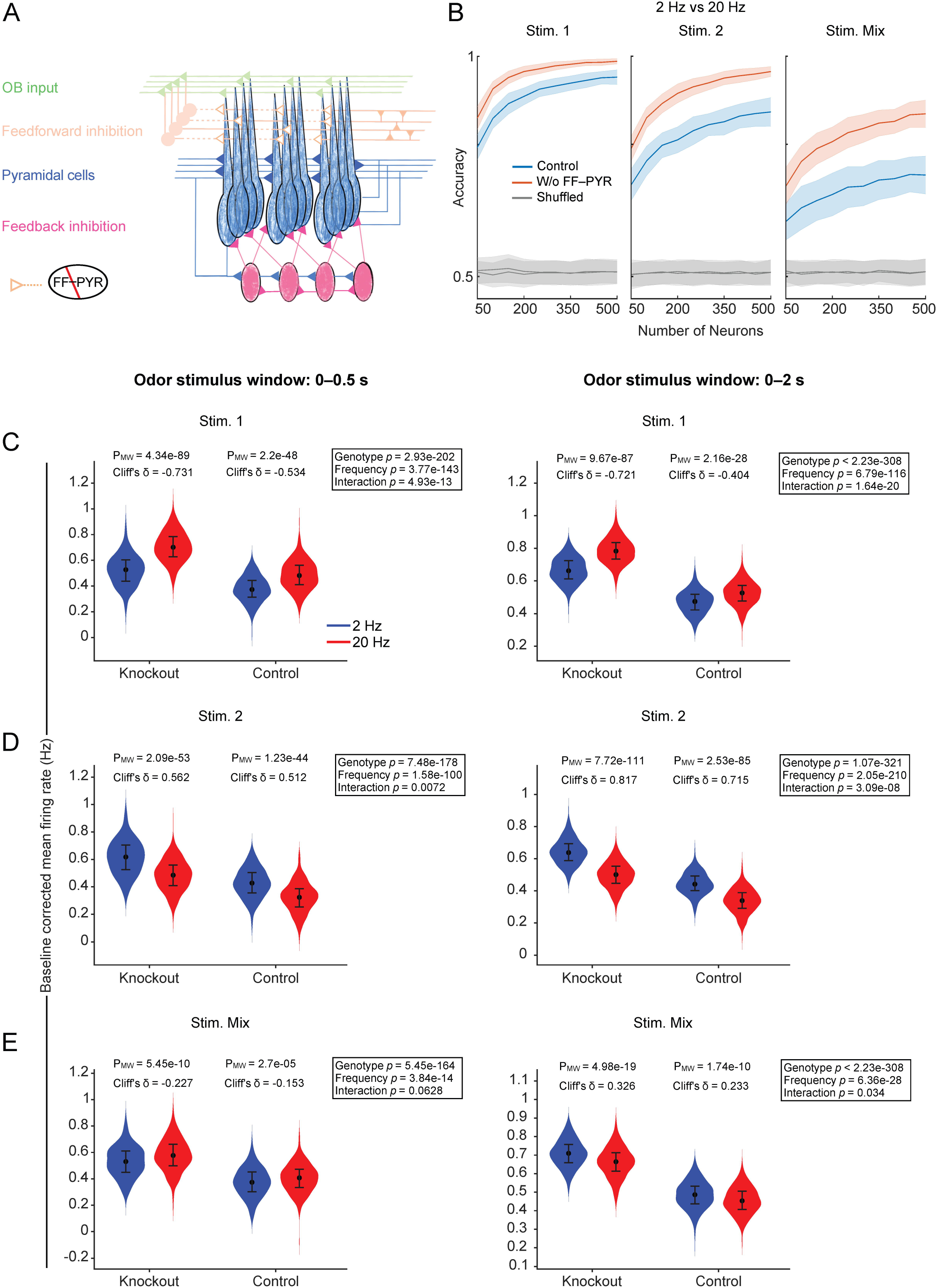
Eliminating the feedforward inhibition (FF) onto PYRs increases the frequency discrimination accuracy of the PCx network. **A,** Schematic of a PCx knockout model in which synaptic connections from feedforward interneurons (FFINs) onto pyramidal neurons (PYRs) were removed. Knocked-out synaptic connections are indicated by hollow triangles. **B,** Average frequency classification accuracy as a function of PYR population size, using outputs from the control (blue) and knockout (red) network models. Solid lines represent the mean, and the shaded regions represent the corresponding standard deviation (*n* = 144). Grey lines represent mean accuracy obtained from the shuffled controls with their corresponding SD. **C,** Violin plots of baseline-corrected mean firing rates (averaged across 336 trials per neuron, *n = 500*) comparing knockout vs. control PCx models in response to 2 Hz and 20 Hz stimulation of stimulus 1; (**D**) stimulus 2; and (**E**) stimulus Mix, during 0–0.5 s (left), and 0–2 s (right) of the stimulus windows. Statistical significance of the 2 Hz vs. 20 Hz comparison was tested using two-tailed Mann–Whitney U tests, and effect sizes were calculated using Cliff’s delta (δ) (*n* = 500 neurons). Two-way mixed repeated measures ART-ANOVA with genotype and frequency as factors.

**Figure 4.**
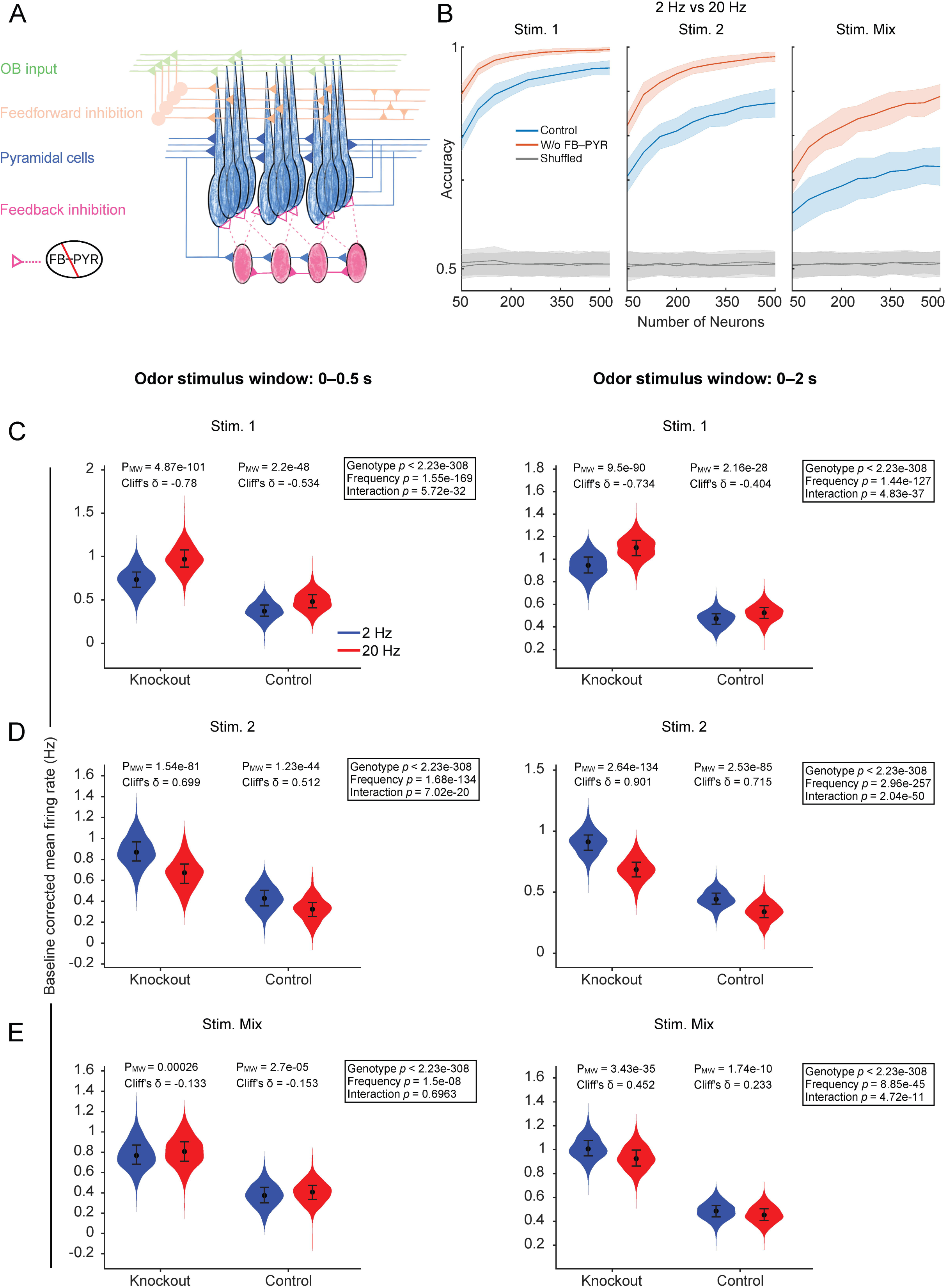
Eliminating feedback inhibition onto PYRs increases the frequency discrimination accuracy of the PCx network model. **A,** Schematic of the PCx knockout model in which synaptic connections from feedback interneurons (FBINs) onto pyramidal neurons (PYRs) were removed. The knocked-out synaptic connections are indicated by hollow triangles. **B,** Odor frequency classification accuracy as a function of PYR population size in control (blue) and knockout (red) PCx using 1D convolutional neural network models. Data are shown as mean ± SD from 144 iterations. Grey lines represent mean accuracy obtained from the shuffled controls with their corresponding SD. **C,** Violin plots of baseline-corrected mean firing rates (averaged across 336 trials per neuron, *n* = 500) comparing knockout vs. control PCx models in response to 2 Hz and 20 Hz stimulation with stimulus 1; (**D**) stimulus 2; and (**E**) stimulus Mix, during 0–0.5 s (left), and 0–2 s (right) of the stimulus windows. Statistical tests were performed as described in Fig. 3C–E.

For stimulus 1, during the 0–0.5 s post-stimulus window, the median firing rate of PYRs increased from 0.53 Hz (IQR = 0.44–0.60 Hz) at 2 Hz stimulation to 0.70 Hz (IQR = 0.63–0.78 Hz) at 20 Hz stimulation (Mann–Whitney *p* = 4.34e−89; Cliff’s |δ| = 0.731, “large” effect size; Fig. 3C, left). We compared the results with the control case as well. Mixed two-way repeated-measures ART-ANOVA revealed significant main effects of genotype (*p* = 2.93e−202) and frequency (*p* = 3.77e−143), as well as a genotype × frequency interaction (*p* = 4.93e−13). A significant genotype × frequency interaction indicates that the two genotypes differ statistically in their degree of change in firing rates across stimulation frequencies. The magnitude of this change is reflected by Cliff’s |δ|, which quantifies the degree of distributional non-overlap between two distributions (47,79). Thus, a genotype demonstrates statistically stronger frequency discrimination when it exhibits a higher Cliff’s |δ| relative to the other along with a significant genotype × frequency interaction.

During the 0–2 s post-stimulus window, the median firing rate increased from 0.66 Hz (IQR = 0.61–0.72 Hz) at 2 Hz to 0.78 Hz (IQR = 0.73–0.84 Hz) at 20 Hz (Mann–Whitney *p* = 9.67e−87; Cliff’s |δ| = 0.721, “large” effect size; Fig. 3C, right). The ART-ANOVA analysis with the corresponding control case again revealed significant main effects of genotype (*p* < 2.225e−308), frequency (*p* = 6.79e−116), and genotype × frequency interaction (*p* = 1.65e−20). Please refer to Table 3 to see the complete summary of the *p* values for the main and interaction effects. Based on this criterion, FF–PYR knockout mice exhibited enhanced odor-frequency discrimination for stimulus 1 across both time windows compared to control.

**Table 3.** . Highlighted *p*-values of contrasts using ART-ANOVA.

| Knockout Type | Effect | Time bin (0–0.5 s) |  |  | Time bin (0–2 s) |  |  |
| --- | --- | --- | --- | --- | --- | --- | --- |
|  |  | Stimulus 1 | Stimulus 2 | Stimulus Mix | Stimulus 1 | Stimulus 2 | Stimulus Mix |
| W/o FF–PYR | Control vs. Knockout | 8.043E–233 | 3.783E–210 | 2.075E–193 | <2.225E–308 | <2.225E–308 | <2.225E–308 |
|  | 2 Hz vs. 20 Hz | 3.290E–160 | 4.600E–108 | 3.957E–14 | 3.094E–125 | 2.727E–241 | 6.362E–28 |
|  | Interaction | 2.173E–13 | 0.007 | 0.064 | 8.782E–21 | 1.934E–08 | 0.035 |
| W/o FB–PYR | Control vs. Knockout | <2.225E–308 | <2.225E–308 | <2.225E–308 | <2.225E–308 | <2.225E–308 | <2.225E–308 |
|  | 2 Hz vs. 20 Hz | 2.406E–186 | 5.566E–147 | 1.531E–08 | 4.979E–139 | 8.252E–299 | 5.914E–47 |
|  | Interaction | 3.175E–32 | 3.899E–20 | 0.696 | 5.523E–38 | 3.951E–52 | 2.648E–11 |
| W/o FF–FF | Control vs. Knockout | 1.847E–39 | 1.606E–22 | 2.104E–25 | 2.680E–87 | 4.795E–55 | 9.072E–82 |
|  | 2 Hz vs. 20 Hz | 1.376E–100 | 1.618E–90 | 1.572E–09 | 6.222E–65 | 2.258E–187 | 3.322E–21 |
|  | Interaction | 0.251 | 0.242 | 0.935 | 0.298 | 0.046 | 0.908 |
| W/o FB–FB | Control vs. Knockout | 3.640E–27 | 8.862E–16 | 1.524E–20 | 9.234E–52 | 1.844E–32 | 8.313E–44 |
|  | 2 Hz vs. 20 Hz | 5.111E–105 | 1.414E–81 | 6.524E–14 | 6.357E–74 | 5.794E–185 | 1.195E–15 |
|  | Interaction | 0.952 | 0.015 | 0.190 | 0.063 | 0.031 | 0.144 |
| W/o FF–FF&FB–FB | Control vs. Knockout | 9.251E–85 | 4.542E–43 | 6.043E–60 | 6.364E–179 | 1.168E–107 | 1.159E–172 |
|  | 2 Hz vs. 20 Hz | 4.904E–82 | 2.701E–66 | 0.0007 | 2.570E–53 | 9.577E–156 | 3.415E–24 |
|  | Interaction | 0.001 | 4.502E–05 | 0.004 | 0.495 | 5.955E–06 | 0.494 |
| W/o PYR–PYR | Control vs. Knockout | 5.786E–63 | 8.046E–37 | 3.543E–61 | 1.954E–142 | 8.882E–95 | 7.569E–161 |
|  | 2 Hz vs. 20 Hz | 1.030E–119 | 1.443E–77 | 9.210E–07 | 1.479E–76 | 3.659E–177 | 1.159E–20 |
|  | Interaction | 0.207 | 0.003 | 0.169 | 0.007 | 0.0007 | 0.872 |

For stimulus 2, during 0–0.5 s post-stimulus window, the median firing rate decreased from 0.62 Hz (IQR = 0.53–0.70 Hz) at 2 Hz to 0.48 Hz (IQR = 0.41–0.56 Hz) at 20 Hz (Mann–Whitney *p* = 2.09e−53; Cliff’s |δ| = 0.562, “large” effect size; Fig. 3D, left). Comparing it with the control, ART-ANOVA revealed significant main effects of genotype (*p* = 7.48e−178), frequency (*p* = 1.58e−100), and a modest genotype × frequency interaction (*p* = 0.007). During the 0–2 s post stimulation window, the median firing rate decreased from 0.64 Hz (IQR = 0.59–0.69 Hz) at 2 Hz to 0.5 Hz (IQR = 0.45–0.55 Hz) at 20 Hz (Mann–Whitney *p* = 7.72e−111; Cliff’s |δ| = 0.817, “large” effect size; Fig. 3D, right). Comparing it with the control, ART-ANOVA again indicated significant main effects of genotype (*p* = 1.07e−321), frequency (*p* = 2.05e−210), as well as genotype × frequency interaction (*p* = 3.08e−08). These results also supported enhanced odor-frequency discrimination for stimulus 2 in FF–PYR knockout compared to control.

For stimulus Mix, during the 0–0.5 s post-stimulus window, median firing rate of PYRs increased modestly from 0.53 Hz (IQR = 0.45–0.61 Hz) at 2 Hz to 0.58 Hz (IQR = 0.50–0.66 Hz) at 20 Hz (Mann–Whitney *p* = 5.45e−10; Cliff’s |δ| = 0.227, “small” effect size; Fig. 3E, left). Comparing it with the control, ART-ANOVA revealed significant main effects of genotype (*p* = 5.45e−164), frequency (*p* = 3.84e−14), but without genotype × frequency interaction (*p* = 0.063). However, during 0–2 s post-stimulus window, the median firing rate decreased from 0.71 Hz (IQR = 0.66–0.76 Hz) at 2 Hz to 0.66 Hz (IQR = 0.61–0.71 Hz) at 20 Hz (Mann–Whitney *p* = 4.98e−19; Cliff’s |δ| = 0.326, “small” effect size; Fig. 3E, right). Comparing it with the control, ART-ANOVA indicated significant effects of genotype (*p* < 2.225e−308), frequency (*p* = 6.36e−28), and a genotype × frequency interaction (*p* = 0.034). These findings indicate improved odor-frequency discrimination in FF–PYR knockout relative to control for stimulus Mix only during the entire 0–2 s window of odor stimulation. Overall, based on the statistical tests, the CNNs and the logistic regression classifiers, FF–PYR knockout model exhibited robust discrimination between 2 Hz and 20 Hz stimulation frequencies relative to control across all odor stimuli and largely across both the analysis windows.

Next, we assessed the odor-frequency discrimination in FB–PYR model. For stimulus 1, the FB–PYR knockout model exhibited stronger discrimination between stimulation frequencies relative to control across both odor-stimulation time windows. During the 0–0.5 s post-stimulus window, median firing rates in the knockout model increased from 0.73 Hz (IQR: 0.64–0.82 Hz) at 2 Hz to 0.97 Hz (IQR: 0.88–1.08 Hz) at 20 Hz (Mann–Whitney *p* = 4.87e−101; Cliff’s |δ| = 0.78, “large” effect size; Fig. 4C, left). Comparing it with the control, ART-ANOVA revealed significant main effects of genotype (*p* < 2.225e−308) and frequency (*p* = 1.55e−169), as well as a genotype × frequency interaction (*p* = 5.72e−32). During the 0–2 s window, median firing rates increased from 0.95 Hz (IQR: 0.88–1.02 Hz) at 2 Hz to 1.10 Hz (IQR: 1.03–1.17 Hz) at 20 Hz (Mann–Whitney *p* = 9.5e−90; Cliff’s |δ| = 0.734, “large” effect size; Fig. 4C, right). Comparing it with control, ART-ANOVA revealed significant main effects of genotype (*p* < 2.225e−308) and frequency (*p* = 1.44e−127), as well as a genotype × frequency interaction (*p* = 4.84e−37).

In case of stimulus 2, the FB–PYR knockout model also exhibited greater frequency discrimination relative to control across both post-stimulation time windows. During the 0–0.5 s post-stimulus window, median firing rates decreased from 0.87 Hz (IQR: 0.78–0.97 Hz) at 2 Hz to 0.67 Hz (IQR: 0.57–0.76 Hz) at 20 Hz (Mann–Whitney *p* = 1.54e−81; Cliff’s |δ| = 0.699, “large” effect size; Fig. 4D, left). Comparing it with control, ART-ANOVA revealed significant main effects of genotype (*p* < 2.225e−308) and frequency (*p* = 1.68e−134), as well as a genotype × frequency interaction (*p* = 7.02e−20). During the 0–2 s post-stimulus window, median firing rates decreased from 0.91 Hz (IQR: 0.84–0.97 Hz) at 2 Hz to 0.68 Hz (IQR: 0.62–0.75 Hz) at 20 Hz (Mann–Whitney *p* = 2.64e−134; Cliff’s |δ| = 0.901, “large” effect size; Fig. 4D, right). Comparing with control, ART-ANOVA revealed significant main effects of genotype (*p* < 2.225e−308) and frequency (*p* = 2.96e−257), as well as a genotype × frequency interaction (*p* = 2.04e−50).

For stimulus Mix, both the FB–PYR knockout and control models showed limited and comparable discrimination between stimulation frequencies during the 0–0.5 s post-stimulus window. Median firing rates in the knockout model increased slightly from 0.77 Hz (IQR: 0.68–0.87 Hz) at 2 Hz stimulation to 0.81 Hz (IQR: 0.71–0.90 Hz) at 20 Hz stimulation (Mann–Whitney *p* = 0.00026; Cliff’s |δ| = 0.133, “negligible” effect size; Fig. 4E, left). Comparing with control, ART-ANOVA revealed significant main effects of genotype (*p* < 2.225e−308), frequency (*p* = 1.5e−08), but no genotype × frequency interaction (*p* = 0.696). However, during the 0–2 s post-stimulus window, median firing rates in the knockout model decreased from 1.01 Hz (IQR: 0.95–1.08 Hz) at 2 Hz stimulation to 0.93 Hz (IQR: 0.86–1.00 Hz) at 20 Hz stimulation (Mann–Whitney *p* = 3.43e−35; Cliff’s |δ| = 0.452, “medium” effect size; Fig. 4E, right). Comparing it with control, ART-ANOVA revealed significant main effects of genotype (*p* < 2.225e−308) and frequency (*p* = 8.85e−45), as well as a genotype × frequency interaction (*p* = 4.72e−11).

These findings are in line with the observations from both linear (logistic regression) and non-linear (1D-CNN) models that both FF–PYR and FB–PYR knockout models exhibited enhanced odor-frequency discrimination across all three odor stimuli over the entire 2 s odor stimulation period. Although direct inhibition appears detrimental to discrimination performance, this must be interpreted in the context of the computational framework. In the present study, added synaptic noise maintained near-physiological baseline firing rates, thereby preventing runaway excitation and enabling the simulation of extreme inhibitory loss. Nevertheless, the perturbation results reveal the network’s capacity, demonstrating that eliminating direct inhibition onto PYRs significantly enhances odor frequency discrimination.

### Eliminating intra-inhibitory connections alone had minimal effect on odor frequency discrimination

The PCx network contains a substantial amount of intra-inhibitory connections among feedforward (FF) as well as feedback (FB) interneurons (43). To understand the influence of these disinhibitory motifs on odor frequency discrimination, we eliminated the synaptic intra-connections among FF and FB interneurons individually in two distinct knockout models (Fig. 5A, 6A). Both knockout models demonstrated comparable decoding accuracy to the control network across various population sizes (Fig. 5B; Fig. 6B; Fig. S6). Peak accuracy achieved by CNNs for the FF–FF knockout model was 0.92 ± 0.016 (stimulus 1), 0.87 ± 0.023 (stimulus 2), and 0.7 ± 0.05 (stimulus Mix), whereas for FB–FB knockout model was 0.93 ± 0.015 (stimulus 1), 0.87 ± 0.028 (stimulus 2), and 0.73 ± 0.04 (stimulus Mix) (Fig. 5B, 6B). Logistic regression classifiers achieved peak decoding accuracies of 0.85 ± 0.019, 0.82 ± 0.018, and 0.68 ± 0.023 for stimuli 1, 2, and Mix, respectively, in the FF–FF knockout model and 0.86 ± 0.019, 0.82 ± 0.02, and 0.71 ± 0.025 for the same stimulus pairs in the FB–FB knockout model (Fig. S6).

**Figure 5.**
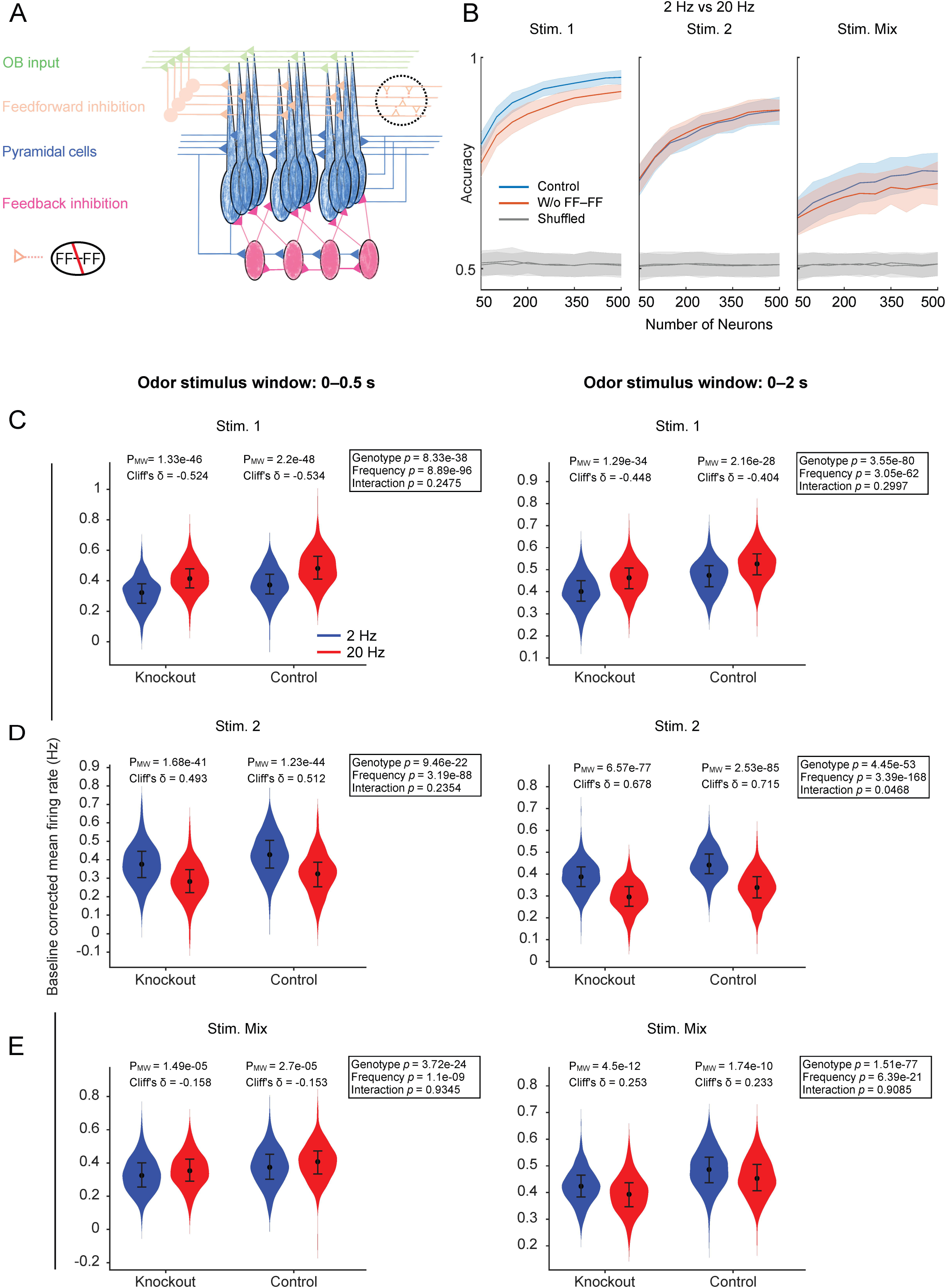
Eliminating recurrency among the FFINs mildly decreased frequency discrimination accuracy of the PCx network model. **A,** Schematic of PCx knockout model in which synaptic connections among feedforward interneurons (FFINs) were removed. Knocked-out synaptic connections are indicated by hollow triangles. **B,** Odor frequency discrimination accuracy as a function of PYR population size in control (blue) and knockout (red) PCx using 1D convolutional neural network models. Data are shown as mean ± SD from 144 iterations. Grey lines represent mean accuracy obtained from the shuffled controls with their corresponding SD. **C,** Violin plots of baseline-corrected mean firing rates (averaged across 336 trials per neuron, *n* = 500) comparing knockout vs. control PCx models in response to 2 Hz and 20 Hz stimulation of stimulus 1; (**D**) stimulus 2; and (**E**) stimulus Mix, during 0–0.5 s (left), and 0–2 s (right) post-stimulus windows. Statistical tests were performed as described in Fig. 3C–E.

**Figure 6.**
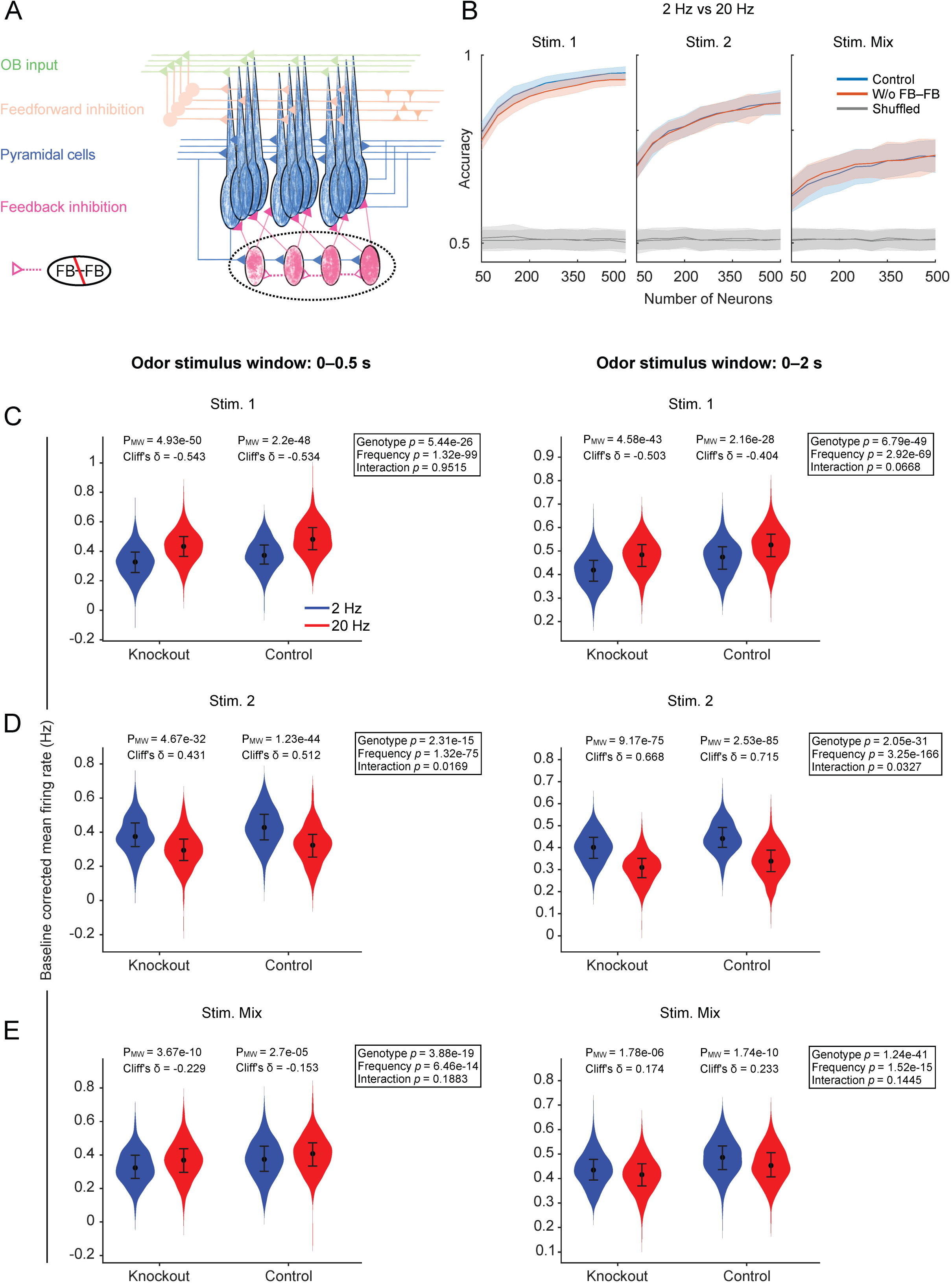
Eliminating recurrency among the FBINs did not alter frequency discrimination accuracy of the PCx network model. **A,** Schematic of the PCx knockout model in which synaptic connections among feedback interneurons (FBINs) were removed. Knocked-out synaptic connections are indicated by hollow triangles. **B,** Odor frequency classification accuracy as a function of PYR population size in control (blue) and knockout (red) PCx using 1D convolutional neural network models. Data are shown as mean ± SD from 144 iterations. Grey lines represent mean accuracy obtained from the shuffled controls with their corresponding SD. **C,** Violin plots of baseline-corrected mean firing rates (averaged across 336 trials per neuron, *n* = 500) comparing knockout vs. control PCx models in response to 2 Hz and 20 Hz stimulation of stimulus 1; (**D**) stimulus 2; and (**E**) stimulus Mix, during 0–0.5 s (left), and 0–2 s (right) post-stimulus windows. Statistical tests were performed as described in Fig. 3C–E.

For stimulus 1, frequency discrimination in the FF–FF knockout model was comparable to that of the control when responses were analyzed during either the initial 0–0.5 s or entire 0–2 s window of odor stimulation (Fig. 5C). During the initial 0–0.5 s window, the median firing rate of PYRs increased from 0.32 Hz (IQR: 0.25–0.38 Hz) at 2 Hz stimulation to 0.41 Hz (IQR: 0.35–0.48 Hz) at 20 Hz stimulation (Mann–Whitney *p* = 1.33e−46; Cliff’s |δ| = 0.524, “large” effect size; Fig. 5C, left). Relative to control, ART-ANOVA showed significant main effects of genotype (*p* = 8.33e−38) and frequency (*p* = 8.89e−96), with no genotype × frequency interaction (*p* = 0.2475). During the 0–2 s window, the median firing rate increased from 0.40 Hz (IQR: 0.36–0.45 Hz) at 2 Hz stimulation to 0.46 Hz (IQR: 0.41–0.51 Hz) at 20 Hz stimulation (Mann–Whitney *p* = 1.29e−34; Cliff’s |δ| = 0.448, “medium” effect size; Fig. 5C, right). ART-ANOVA again showed significant main effects of genotype (*p* = 3.55e−80) and frequency (*p* = 3.05e−62), with no genotype × frequency interaction (*p* = 0.2997).

For stimulus 2, frequency discrimination remained comparable to control during the 0–0.5 s window. Here PYRs’ median firing rate decreased from 0.38 Hz (IQR: 0.30–0.45 Hz) at 2 Hz stimulation to 0.28 Hz (IQR: 0.22–0.35 Hz) at 20 Hz stimulation (Mann–Whitney *p* = 1.68e−41; Cliff’s |δ| = 0.493, “large” effect size; Fig. 5D, left). ART-ANOVA revealed significant main effects of genotype (*p* = 9.46e−22) and frequency (*p* = 3.19e−88), with no genotype × frequency interaction (*p* = 0.2354). During the 0–2 s window, PYRs’ median firing rate decreased from 0.39 Hz (IQR: 0.34–0.43 Hz) at 2 Hz stimulation to 0.30 Hz (IQR: 0.25–0.34 Hz) at 20 Hz stimulation (Mann–Whitney *p* = 6.57e−77; Cliff’s |δ| = 0.678, “large” effect size; Fig. 5D, right). However, for this time window, ART-ANOVA revealed a significant genotype × frequency interaction (*p* = 0.0468), alongside significant main effects of genotype (*p* = 4.45e−53) and frequency (*p* = 3.39e−168), indicating slightly higher discrimination in control.

For the stimulus Mix, frequency discrimination in the knockout model remained comparable to control for both time windows (Fig. 5E). During the 0–0.5 s window, the median firing rate of PYRs increased slightly from 0.32 Hz (IQR: 0.25–0.40 Hz) at 2 Hz stimulation to 0.35 Hz (IQR: 0.29–0.42 Hz) at 20 Hz stimulation (Mann–Whitney *p* = 1.49e−05; Cliff’s |δ| = 0.158, “small” effect size; Fig. 5E, left). ART-ANOVA showed significant main effects of genotype (*p* = 3.72e−24) and frequency (*p* = 1.1e−09), with no genotype × frequency interaction (*p* = 0.9345). During the 0–2 s window, the median firing rate decreased from 0.42 Hz (IQR: 0.38–0.47 Hz) at 2 Hz stimulation to 0.39 Hz (IQR: 0.35–0.44 Hz) at 20 Hz stimulation (Mann–Whitney *p* = 4.5e−12; Cliff’s |δ| = 0.253, “small” effect size; Fig. 5E, right). ART-ANOVA again revealed significant main effects of genotype (*p* = 1.51e−77) and frequency (*p* = 6.39e−21) without a genotype × frequency interaction (*p* = 0.9085).

Next, the control and FB–FB knockout models both exhibited comparable odor frequency discrimination for stimuli 1 and Mix (Fig. 6C,E). However, for stimulus 2, the knockout model exhibited reduced discrimination ability compared to the control across both stimulus time windows (Fig. 6D). For stimulus 1, during the 0–0.5 s window, PYRs’ median firing rate increased substantially from 0.33 Hz (IQR: 0.25–0.39 Hz) at 2 Hz stimulation to 0.43 Hz (IQR: 0.36–0.50 Hz) at 20 Hz stimulation (Mann–Whitney *p* = 4.93e−50; Cliff’s |δ| = 0.543, “large” effect size; Fig. 6C, left). ART-ANOVA showed significant main effects of genotype (*p* = 5.44e−26) and frequency (*p* = 1.32e−99), with no genotype × frequency interaction (*p* = 0.9515). During the 0–2 s window, median firing rate increased from 0.42 Hz (IQR: 0.37–0.46 Hz) at 2 Hz stimulation to 0.48 Hz (IQR: 0.43–0.53 Hz) at 20 Hz stimulation (Mann–Whitney *p* = 4.58e−43; Cliff’s |δ| = 0.503, “large” effect size; Fig. 6C, right). ART-ANOVA again revealed significant main effects of genotype (*p* = 6.79e−49) and frequency (*p* = 2.92e−69) without a genotype × frequency interaction (*p* = 0.0668).

For stimulus 2, during the 0–0.5 s window, median firing rate decreased from 0.37 Hz (IQR: 0.32–0.45 Hz) at 2 Hz stimulation to 0.29 Hz (IQR: 0.23–0.36 Hz) at 20 Hz stimulation (Mann–Whitney *p* = 4.67e−32; Cliff’s |δ| = 0.431, “medium” effect size; Fig. 6D, left). The higher Cliff’s |δ| in control relative to knockout, along with significant main effects of genotype (*p* = 2.31e−15), frequency (*p* = 1.32e−75), and a genotype × frequency interaction (*p* = 0.0169) by ART-ANOVA indicated better discrimination in control. During the 0–2 s window, PYRs’ median firing rate decreased markedly from 0.40 Hz (IQR: 0.35–0.45 Hz) at 2 Hz stimulation to 0.31 Hz (IQR: 0.26–0.35 Hz) at 20 Hz stimulation (Mann–Whitney *p* = 9.17e−75; Cliff’s |δ| = 0.668, “large” effect size; Fig. 6D, right). The lower Cliff’s |δ| in knockout relative to control, along with significant main effects of genotype (*p* = 2.05e−31), frequency (*p* = 3.25e−166), and a genotype × frequency interaction (*p* = 0.0327), confirmed marginally impaired discrimination in the knockout model.

For the stimulus Mix, frequency discrimination in the knockout model remained comparable to control across both time windows (Fig. 6E). During the 0–0.5 s window, the median firing rate of PYRs increased modestly from 0.32 Hz (IQR: 0.26–0.40 Hz) at 2 Hz stimulation to 0.37 Hz (IQR: 0.30–0.44 Hz) at 20 Hz stimulation (Mann–Whitney *p* = 3.67e−10; Cliff’s |δ| = 0.229, “small” effect size; Fig. 6E, left). ART-ANOVA showed significant main effects of genotype (*p* = 3.88e−19) and frequency (*p* = 6.46e−14), with no genotype × frequency interaction (*p* = 0.1883). During the 0–2 s window, the median firing rate showed a minimal decrease from 0.43 Hz (IQR: 0.39–0.48 Hz) at 2 Hz stimulation to 0.42 Hz (IQR: 0.37–0.46 Hz) at 20 Hz stimulation (Mann–Whitney *p* = 1.78e−06; Cliff’s |δ| = 0.174, “small” effect size; Fig. 6E, right). ART-ANOVA again revealed significant main effects of genotype (*p* = 1.24e−41) and frequency (*p* = 1.52e−15) without a genotype × frequency interaction (*p* = 0.1445).

Overall, our results demonstrate that eliminating the intra-inhibitory connections within either feedforward or feedback interneuron populations did not substantially affect the odor frequency discrimination capacity of the PCx.

### Eliminating either excitatory or all intra-inhibitory recurrent connections impaired odor frequency discrimination

Finally, to investigate how the simultaneous removal of recurrent inhibition in both interneuron types affects frequency encoding, we developed a model that exhibited both FF–FF and FB–FB knockouts (Fig. 7A). To investigate the effect of removing recurrent excitation onto PYRs, we developed a PYR–PYR knockout model (Fig. 8A). Decoding of odor frequency responses using 1D-CNN models revealed a decline in classification accuracy for both knockout models across all size-matched populations of PYRs compared to the control (Fig. 7B, 8B). The CNN models achieved peak decoding accuracies of 0.91 ± 0.02, 0.83 ± 0.029, and 0.67 ± 0.05 for stimuli 1, 2, and Mix, respectively, in the dual intra-inhibitory knockout model (Fig. 7B). The peak accuracies achieved in the PYR–PYR knockout model were 0.92 ± 0.02, 0.84 ± 0.03, and 0.69 ± 0.05 for stimuli 1, 2, and Mix respectively (Fig. 8B) compared to the control (stimulus 1: 0.95 ± 0.016; stimulus 2: 0.87 ± 0.032; and Mix: 0.73 ± 0.043).

**Figure 7.**
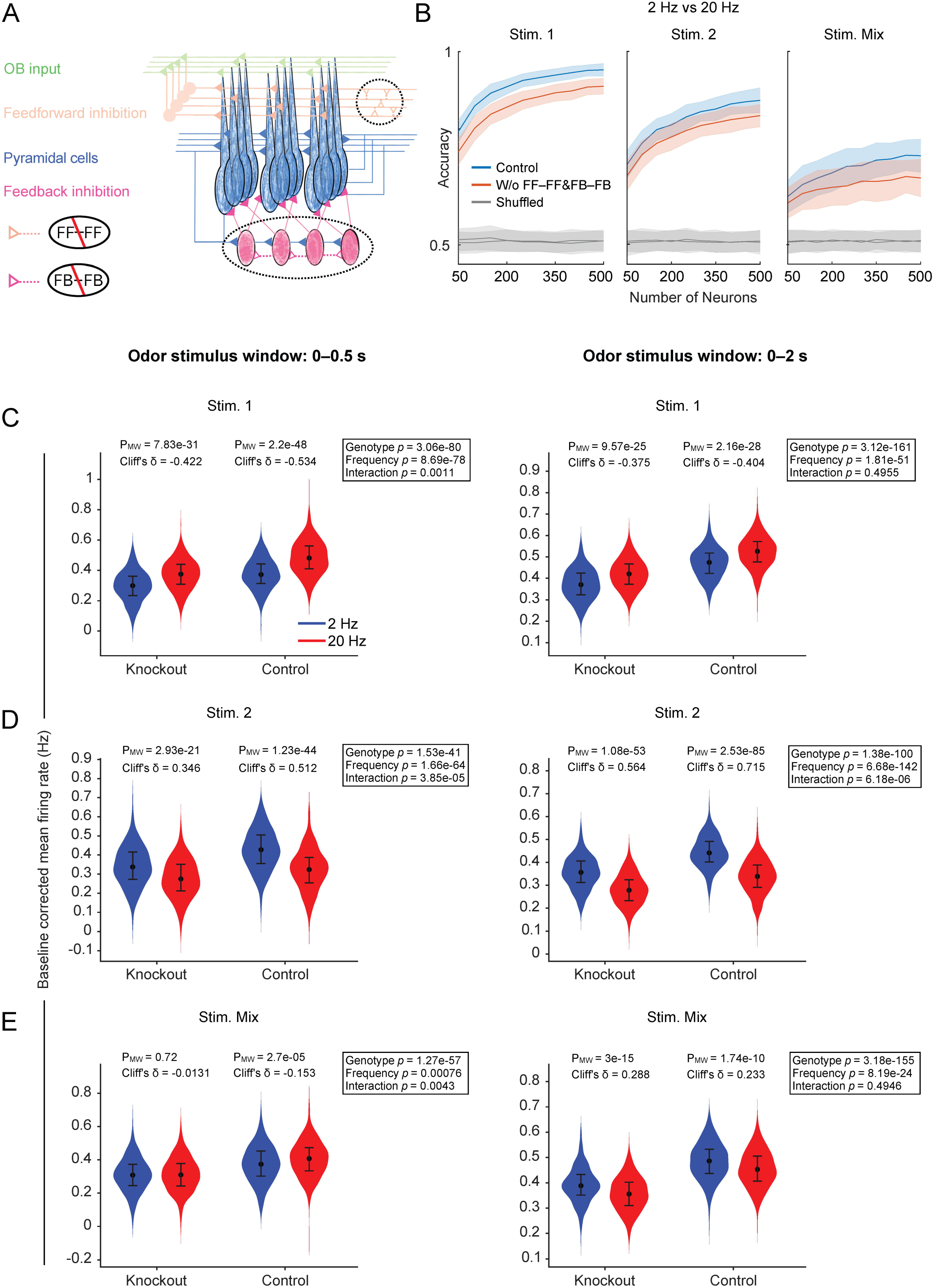
Impairment of frequency discrimination accuracy of the PCx network model due to the elimination of recurrent inhibition within both FFINs and FBINs. **A,** Schematic of the PCx knockout model in which synaptic connections among both feedforward interneurons (FFINs) and feedback interneurons (FBINs) were removed. Knocked-out synaptic connections are indicated by hollow triangles. **B,** Odor frequency classification accuracy as a function of PYR population size in control (blue) and knockout (red) PCx using 1D convolutional neural network models. Data are shown as mean ± SD from 144 iterations. Grey lines represent mean accuracy obtained from the shuffled controls with their corresponding SD. **C,** Violin plots of baseline-corrected mean firing rates (averaged across 336 trials per neuron, *n* = 500) comparing knockout vs. control PCx models in response to 2 Hz and 20 Hz stimulation of stimulus 1; (**D**) stimulus 2; and (**E**) stimulus Mix, during 0–0.5 s (left), and 0–2 s (right) post-stimulus windows. Statistical tests were performed as described in Fig. 3C–E.

**Figure 8.**
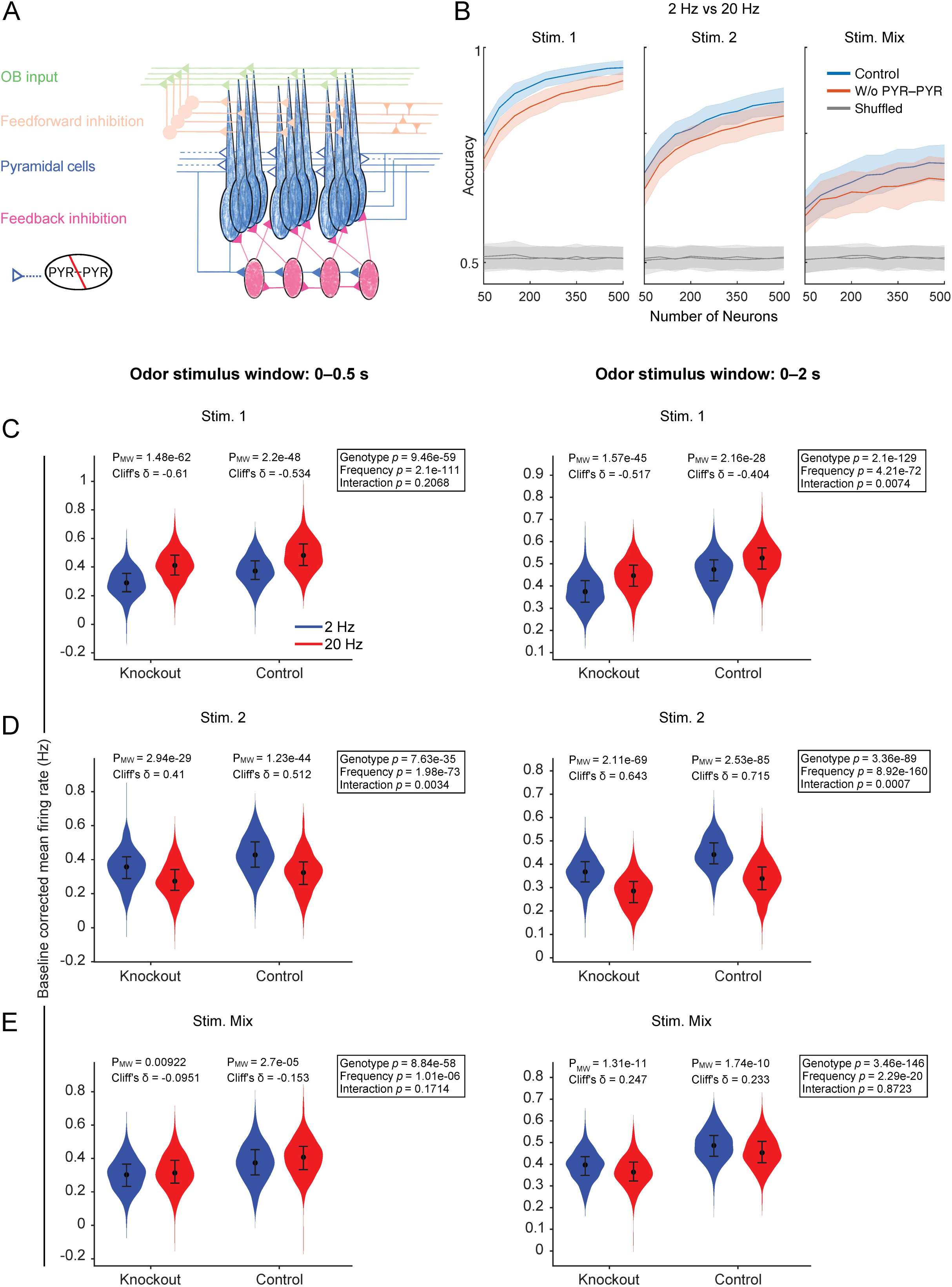
Eliminating recurrency among the PYRs decreased the frequency discrimination accuracy of the PCx network model. **A,** Schematic of the PCx knockout model in which recurrent synaptic connections among pyramidal neurons (PYRs) were removed. Knocked-out synaptic connections are indicated by hollow triangles. **B,** Odor frequency classification accuracy as a function of PYR population size in control (blue) and knockout (red) PCx using 1D convolutional neural network models. Data are shown as mean ± SD from 144 iterations. Grey lines represent mean accuracy obtained from the shuffled controls with their corresponding SD. **C,** Violin plots of baseline-corrected mean firing rates (averaged across 336 trials per neuron, *n* = 500) comparing knockout vs. control PCx models in response to 2 Hz and 20 Hz stimulation of stimulus 1; (**D**) stimulus 2; and (**E**) stimulus Mix, during 0–0.5 s (left), and 0–2 s (right) post-stimulus windows. Statistical tests were performed as described in Fig. 3C–E.

**Figure 9.**
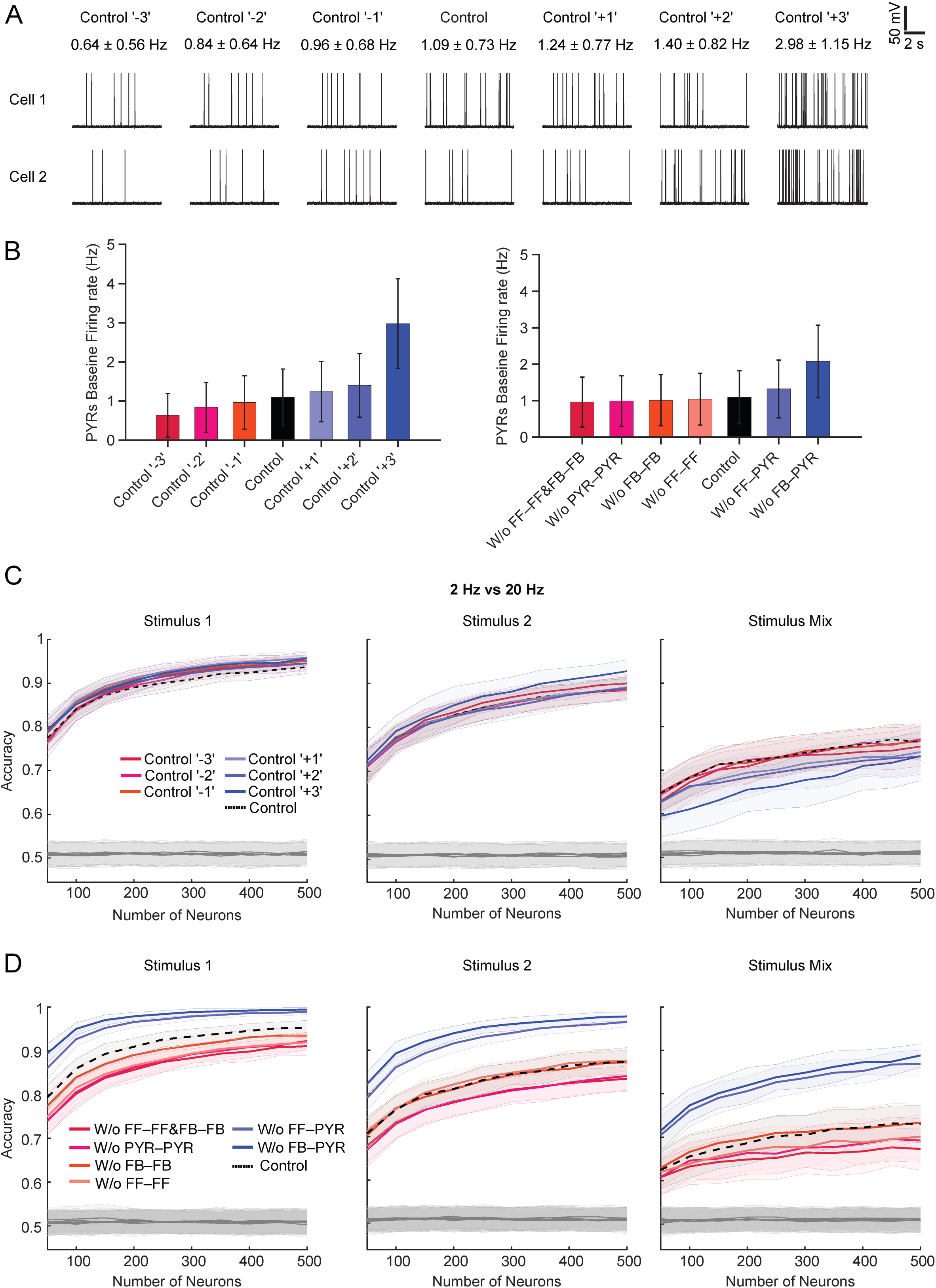
Baseline firing rate does not determine odor frequency discrimination in the PCx. **A,** Example voltage traces from PYRs in control and variant networks (Control ‘-3’ to Control ‘+3’), showing baseline activity levels modulated by varying the baseline noise, resulting in firing rates ranging from 0.635 ± 0.56 Hz to 2.98 ± 1.14 Hz (left to right). **B,** Baseline firing rates of the PYR population across control variants (left panel) and knockout models (right panel). Bar heights represent mean firing rates, and error bars indicate standard deviations (*n* = 336 trials). **C,** Odor frequency discrimination accuracy using 1D-CNNs as a function of PYR population size for control variants across three odor stimuli conditions (stimulus 1, stimulus 2, and stimulus Mix). Solid lines represents the mean, and the shaded region represents the standard deviation (*n* = 144 iterations). **D,** Odor frequency decoding accuracy as a function of PYR population size for knockout models across the same three odor stimuli conditions. Solid lines and shaded regions are defined as in Fig. 9C (*n* = 144 iterations).

Logistic regression classifier also displayed similar impairments in decoding accuracies (Fig. S6). The peak decoding accuracies achieved using the responses of FF–FF and FB–FB knockout model were stimulus 1: 0.85 ± 0.02; stimulus 2: 0.78 ± 0.021; and stimulus Mix: 0.64 ± 0.026, whereas those from the PYR–PYR knockout model were stimulus 1: 0.82 ± 0.017; stimulus 2: 0.79 ± 0.02; and Mix: 0.66 ± 0.023 (Fig. S6).

The FF–FF and FB–FB knockout model demonstrated impaired odor frequency discrimination relative to the control for stimuli 1 and Mix during the 0–0.5 s stimulus window, and for stimulus 2 even during the 0–2 s window (Fig. 7C–E). For stimulus 1, during the 0–0.5 s post-stimulus window, PYRs’ median firing rate in the knockout model increased from 0.30 Hz (IQR: 0.23–0.36 Hz) at 2 Hz to 0.37 Hz (IQR: 0.31–0.44 Hz) at 20 Hz (Mann–Whitney *p* = 7.83e−31; Cliff’s |δ| = 0.422, “medium” effect size; Fig. 7C, left). Compared with the control (Cliff’s |δ| = 0.534), ART-ANOVA revealed significant main effects of genotype (*p* = 3.06e−80) and frequency (*p* = 8.69e−78), as well as a genotype × frequency interaction (*p* = 0.0011), indicating impaired frequency discrimination in knockout model. During the 0–2 s window, median firing rate increased from 0.37 Hz (IQR: 0.32–0.42 Hz) at 2 Hz to 0.42 Hz (IQR: 0.37–0.47 Hz) at 20 Hz (Mann–Whitney *p* = 9.57e−25; Cliff’s |δ| = 0.375, “medium” effect size; Fig. 7C, right). Comparing it with control, ART-ANOVA showed significant main effects of genotype (*p* = 3.12e−161) and frequency (*p* = 1.81e−51) but no genotype × frequency interaction (*p* = 0.4955), suggesting comparable discrimination between control and knockout.

For stimulus 2, during the 0–0.5 s window, median firing rate decreased from 0.34 Hz (IQR: 0.27–0.42 Hz) at 2 Hz stimulation to 0.27 Hz (IQR: 0.21–0.35 Hz) at 20 Hz stimulation (Mann–Whitney *p* = 2.93e−21; Cliff’s |δ| = 0.346, “medium” effect size; Fig. 7D, left). The larger effect size observed in control (Cliff’s |δ| = 0.512) relative to knockout, along with significant main effects of genotype (*p* = 1.53e−41), frequency (*p* = 1.66e−64), and a genotype × frequency interaction (*p* = 3.85e−05) by ART-ANOVA, indicated superior discrimination in control. During the 0–2 s window, PYRs median firing rate decreased markedly from 0.36 Hz (IQR: 0.31–0.41 Hz) at 2 Hz stimulation to 0.28 Hz (IQR: 0.23–0.32 Hz) at 20 Hz stimulation (Mann–Whitney *p* = 1.08e−53; Cliff’s |δ| = 0.564, “large” effect size; Fig. 7D, right). The smaller effect size in knockout relative to control (Cliff’s |δ| = 0.715), together with significant main effects of genotype (*p* = 1.38e−100), frequency (*p* = 6.68e−142), and a genotype × frequency interaction (*p* = 6.18e−06), confirmed a significantly impaired discrimination in the knockout model.

During the 0–0.5 s window, there was no significant change in PYRs’ median firing rate in response to 2 Hz vs. 20 Hz stimulation of stimulus Mix in the knockout model (2 Hz stim.: 0.31 Hz, IQR: 0.25–0.37 Hz; 20 Hz stim.: 0.31 Hz, IQR: 0.24–0.38 Hz; Mann–Whitney *p* = 0.72; Cliff’s |δ| = 0.0131, “negligible” effect size; Fig. 7E, left). In contrast, firing rate in control varied significantly across stimulation frequencies (Mann–Whitney *p* = 2.7e−05; Cliff’s |δ| = 0.153; two-way mixed RM ART-ANOVA: Control vs. Knockout, *p* = 1.27e−57; 2 Hz vs. 20 Hz stimulation, *p* = 0.00076; and interaction, *p* = 0.0043), demonstrating substantially weaker discrimination in the knockout model. During the 0–2 s window, changes in PYRs firing rate in knockout model (2 Hz stim.: 0.39 Hz, IQR: 0.35–0.43 Hz; 20 Hz stim.: 0.36 Hz, IQR: 0.31–0.40 Hz; Mann–Whitney *p* = 3e−15; Cliff’s |δ| = 0.288, “small” effect size; Fig. 7E, right) was similar to that observed in control (Control vs. Knockout, *p* = 3.18e−155; 2 Hz vs. 20 Hz stimulation, *p* = 8.19e−24; and interaction, *p* = 0.4946; two-way mixed RM ART-ANOVA).

Next, upon the removal of reciprocal connections between pyramidal neurons, the PYR–PYR knockout model exhibited enhanced discrimination relative to the control for stimulus 1 during 0–2 s time window. Conversely, discrimination was compromised in the case of stimulus 2 across both the time windows, while for stimulus Mix, it remained comparable to control.

For stimulus 1, during the 0–0.5 s window, the knockout model strongly discriminated between the odor frequencies (2 Hz stim.: 0.29 Hz, IQR: 0.23–0.36 Hz; 20 Hz stim.: 0.41 Hz, IQR: 0.34–0.48 Hz; Mann–Whitney *p* = 1.48e−62; Cliff’s |δ| = 0.61, “large” effect size; Fig. 8C, left). A two-way mixed repeated-measures ART-ANOVA revealed main effects of genotype (*p* = 9.46e−59) and frequency (*p* = 2.1e−111) but not interaction (*p* = 0.2068), suggesting that discrimination was similar to control. However, during the 0–2 s window, the knockout model exhibited enhanced discrimination (2 Hz stim.: 0.37 Hz, IQR: 0.33–0.42 Hz; 20 Hz stim.: 0.45 Hz, IQR: 0.40–0.49 Hz; Mann–Whitney *p* = 1.57e−45; Cliff’s |δ| = 0.517, “large” effect size; Fig. 8C, right) relative to control (Cliff’s |δ| = 0.404) as evident by significant main and interaction effects by ART-ANOVA (genotype, *p* = 2.1e−129; frequency, *p* = 4.21e−72; interaction, *p* = 0.0074).

For stimulus 2, during the 0–0.5 s window, the knockout model showed impaired discrimination (2 Hz stim.: 0.36 Hz, IQR: 0.29–0.42 Hz; 20 Hz stim.: 0.27 Hz, IQR: 0.22–0.34 Hz; Mann–Whitney *p* = 2.94e−29; Cliff’s |δ| = 0.41, “medium” effect size; Fig. 8D, left) relative to control (Cliff’s |δ| = 0.512), as explained by significant main and interaction effects by ART-ANOVA (genotype, *p* = 7.63e−35; frequency, *p* = 1.98e−73; interaction, *p* = 0.0034). Similarly, during the 0–2 s window, the knockout model again showed poor discrimination (2 Hz stim.: 0.37 Hz, IQR: 0.32–0.41 Hz; 20 Hz stim.: 0.29 Hz, IQR: 0.24–0.33 Hz; Mann–Whitney *p* = 2.11e−69; Cliff’s |δ| = 0.643, “large” effect size; Fig. 8D, right) relative to control (Cliff’s |δ| = 0.715), as evident by significant main and interaction effects by ART-ANOVA (genotype, *p* = 3.36e−89; frequency, *p* = 8.92e−160; interaction, *p* = 0.0007).

For stimulus Mix, during the 0–0.5 s window, the knockout model was unable to discriminate between two frequencies (2 Hz stim.: 0.30 Hz, IQR: 0.23–0.37 Hz; 20 Hz stim.: 0.31 Hz, IQR: 0.25–0.39 Hz; Mann–Whitney *p* = 0.00922; Cliff’s |δ| = 0.0951, “negligible” effect size; Fig. 8E, left) similar to control (ART-ANOVA: genotype, *p* = 8.84e−58, frequency, *p* = 1.01e−06; and interaction, *p* = 0.1714).

During the 0–2 s window, the knockout model exhibited similar discrimination (2 Hz stim.: 0.40 Hz, IQR: 0.35–0.43 Hz; 20 Hz stim.: 0.36 Hz, IQR: 0.32–0.41 Hz; Mann–Whitney *p* = 1.31e−11; Cliff’s |δ| = 0.247, “small” effect size; Fig. 8E, right) as control (ART-ANOVA: genotype, *p* = 3.46e−146; frequency, *p* = 2.29e−20; interaction, *p* = 0.8723).

In summary, virtual knockout experiments demonstrated that eliminating direct inhibition onto PYRs enhanced frequency discrimination, whereas removing reciprocal connections among either pyramidal neurons, or simultaneously within both interneuron populations degraded decoding performance. Figure S7 and Table 4 summarize the areas under the ROC curves (AUCs) for CNN and logistic regression classifiers to evaluate their performances across control and knockout conditions.

**Table 4.** AUC values of ROC curves.

| Network Model | 1D-CNN |  |  | Logistic Regression Classifier |  |  |
| --- | --- | --- | --- | --- | --- | --- |
|  | Stim. 1 | Stim. 2 | Stim. Mix | Stim. 1 | Stim. 2 | Stim. Mix |
| Control | $0.992 \pm 0.003$ | $0.953 \pm 0.018$ | $0.813 \pm 0.044$ | $0.895 \pm 0.018$ | $0.846 \pm 0.020$ | $0.720 \pm 0.024$ |
| W/o FF-PYR | $0.999 \pm 0.001$ | $0.995 \pm 0.003$ | $0.946 \pm 0.019$ | $0.971 \pm 0.009$ | $0.945 \pm 0.013$ | $0.814 \pm 0.023$ |
| W/o FB-PYR | 1.0 | $0.998 \pm 0.002$ | $0.958 \pm 0.020$ | $0.987 \pm 0.006$ | $0.976 \pm 0.009$ | $0.888 \pm 0.018$ |
| W/o FF-FF | $0.978 \pm 0.008$ | $0.954 \pm 0.015$ | $0.781 \pm 0.042$ | $0.865 \pm 0.017$ | $0.832 \pm 0.021$ | $0.690 \pm 0.026$ |
| W/o FB-FB | $0.986 \pm 0.006$ | $0.949 \pm 0.017$ | $0.822 \pm 0.042$ | $0.865 \pm 0.018$ | $0.832 \pm 0.020$ | $0.715 \pm 0.027$ |
| W/o FF-FF & FB-FB | $0.973 \pm 0.010$ | $0.914 \pm 0.022$ | $0.749 \pm 0.054$ | $0.860 \pm 0.019$ | $0.791 \pm 0.021$ | $0.648 \pm 0.025$ |
| W/o PYR-PYR | $0.977 \pm 0.010$ | $0.922 \pm 0.026$ | $0.768 \pm 0.053$ | $0.823 \pm 0.020$ | $0.798 \pm 0.022$ | $0.664 \pm 0.027$ |

Values are means ± SD; $n = 144$ . W/o, without; PYR, pyramidal neuron; FFIN, feedforward interneuron; FBIN, feedback interneuron.

### Circuit motifs, not just baseline activity level, determine odor frequency representation

Synaptic modifications altered the baseline firing rates of PYRs. Knocking out the direct inhibition onto the PYRs increased baseline firing from 1.09 ± 0.73 Hz (Control) to 1.32 ± 0.79 Hz (FF–PYR knockout) and 2.08 ± 0.99 Hz (FB–PYR knockout) (Fig. 9B). In contrast, knocking out recurrent synaptic connections among PYRs (PYR–PYR knockout) and within both interneuron populations (FF–FF and FB–FB knockout) decreased baseline firing rates to 0.99 ± 0.69 Hz and 0.96 ± 0.68 Hz, respectively. Since changes in odor frequency decoding accuracy correlated with baseline firing rate, we tested whether excitability alone could account for decoding performance. To address this, we simulated control variants of the PCx network by systematically varying the spontaneous firing rates of PYRs (Fig. 9A). The mean spontaneous activity of the PYR population across these variants spanned the entire range observed across the knockout models, from the lower to upper extremes of the firing rate distribution (Fig. 9B). A complete comparative summary of baseline firing rates for each network model is provided in Table 5. Odor frequency decoding was then performed using PYR population responses from control variants (Fig. 9C) and compared with those from the knockout models (Fig. 9D) across all odor conditions. CNN decoding performance using knockout models’ activity revealed a correlation between enhancement or impairment of odor frequency classification and mean population baseline firing rate relative to the control (Fig. 9D). In contrast, control variants showed either minimal differences among classification performances for stimuli 1 and 2, or a reversed correlation for stimulus Mix (Fig. 9C).

**Table 5.** Pyramidal population’s baseline firing rate (Hz)

| W/o<br>FF-FF & FB-FB | W/o<br>PYR-PYR | W/o<br>FB-FB | W/o<br>FF-FF | Control | W/o<br>FF-PYR | W/o<br>FB-PYR |
| --- | --- | --- | --- | --- | --- | --- |
| $0.963 \pm 0.683$ | $0.992 \pm 0.692$ | $1.01 \pm 0.699$ | $1.04 \pm 0.711$ | $1.09 \pm 0.726$ | $1.32 \pm 0.792$ | $2.08 \pm 0.991$ |
| Control '-3' | Control '-2' | Control '-1' | Control | Control '+1' | Control '+2' | Control '+3' |
| $0.635 \pm 0.561$ | $0.839 \pm 0.641$ | $0.961 \pm 0.683$ | $1.09 \pm 0.726$ | $1.24 \pm 0.771$ | $1.4 \pm 0.815$ | $2.98 \pm 1.146$ |

Values are mean ± SD; $n = 336$ iterations for each odor condition across 500 neurons.

Overall, our analysis indicates that specific intracortical circuit motifs in the PCx, rather than the baseline excitability of the PYRs *per se*, are critical for shaping odor frequency representation and encoding.

## Discussion

Mammalian and insect olfactory systems have long been recognized for their ability to encode chemical and physical properties of odor stimuli, such as odor identity, intensity, and concentration (80). These systems also exhibit a remarkable capacity to encode temporal structure of olfactory cues present in natural odor landscapes to guide behavioral decisions in everyday ethological activities such as foraging, navigation, and locating odor sources (1,3–5,7,81). A recent study in mice showed that mitral/tufted (M/T) cells in the OB can encode odor frequencies up to 20 Hz (18). However, whether piriform circuitry can preserve and transform this temporal information remains unresolved. In this study, we investigated whether the PCx can encode odor stimulus frequencies. To this end, we used previously recorded *in vivo* activities from M/Ts in response to odor stimuli at 2 Hz and 20 Hz (18) as input to a biophysically relevant PCx network model. We found that the PCx network model has the capacity to differentially encode 2 Hz and 20 Hz odor stimuli.

We then simulated several synaptic knockout variants of the PCx model to assess how frequency encoding is affected by circuit perturbations, hence interrogating their significance. We found that the odor frequency discrimination of the PCx model was quite robust to changes in the inhibitory strength resulting from the removal of disinhibition in either feedforward or feedback microcircuits. It is possible that the PCx’s discriminative capacity was buffered against the moderate increase in feedforward or feedback inhibition following the removal of mutual inhibition within these microcircuits (43). However, simultaneous removal of disinhibition in both these microcircuits caused a minor but significant impairment in the decoding performance.

The recurrent circuitry of the PCx is known to play a crucial role in the process of pattern completion, attractor dynamics, and stable odor representations of degraded or noisy olfactory inputs (24,25,82–86). By disrupting the recurrent excitation of the pyramidal neurons, we discovered that odor frequency discrimination was significantly deteriorated, consistent with the crucial role of recurrently connected piriform neurons in stabilizing sensory odor representations (22). In contrast, selectively eliminating the direct synaptic inhibition onto pyramidal neurons from feedforward or feedback microcircuits markedly improved odor frequency discrimination. To further investigate the relationship between E/I balance and discrimination performance, different variants of the control model were simulated by enhancing or suppressing PYRs’ baseline activity (Fig. 9C). However, we did not find any direct correlation between spontaneous firing rate and frequency discrimination. These results support the idea that the E/I balance enforced by different microcircuits modulates the frequency encoding in a particular way, which cannot be fully explained by merely changing the baseline excitability of PYRs.

In conclusion, our results indicate that the PCx network can bidirectionally modulate the odor frequency discrimination, a mechanism that is sculpted by various canonical circuit motifs.

### Single-cell vs. population coding

We initially sought to understand how individual pyramidal neurons represent odor-frequency. To investigate this, we employed a cellular-level statistical approach in which individual pyramidal neurons were classified as frequency-discriminating or non-discriminating types based on their odor-evoked firing rates across 336 trials (Mann–Whitney U test; Bonferroni correction) (Fig. 1I–K). We found that individual pyramidal neurons are insufficient to reliably encode odor stimulation frequency. Moreover, such single-neuron encoding would result in a low-dimensional representation of stimulus parameters such as odor frequency, in a systematically patterned network (87). Prior studies, however, have shown that multiple neurons often respond nonlinearly to collectively represent a given stimulus parameter, which in turn helps the organism to perceive it (88–90). Overall, single-neuron analysis is only partially predictive of a cell’s contribution to representing a given stimulus feature (91).

We therefore turned our attention to the population-level decoding of frequency information. The collective activity of all PYRs, quantified using the trial-averaged firing rates of individual neurons across 336 trials, effectively discriminated between the two stimulation frequencies (Fig. 1F–H). However, within a network, the activities of individual neurons are temporally correlated with one another. Stefanini et al. (2020) demonstrated that such correlations may cause a decoder to assign larger weights to the weakly tuned or even untuned neurons when their activities are correlated with tuned neurons. Population decoders (pattern classifiers) using the average firing rate of neurons have been extensively used in different fields of neuroscience to predict decision boundaries from experimental outcomes. However, averaging spiking activity over extended periods results in the loss of valuable information embedded in the temporal structure of the population firing patterns. To preserve this rich temporal information, we employed one-dimensional convolutional neural networks (1D-CNNs), and logistic regression classifiers for odor frequency classification. These models used binned spiking activity of neurons across the entire stimulation period as input, using 20 ms non-overlapping time windows. Each bin represented a 20 ms time interval and was assigned a binary value (0 or 1) indicating whether the neuron fired within that interval. The 20 ms bin size was selected to minimize the probability of capturing more than one spike per bin. Both nonlinear and linear models, trained and tested on PYR population activity, reliably predicted odor stimulus frequency (Fig. 2B,D; S6; S7).

Using the trial-averaged (*n* = 300) log-likelihood difference (ΔLL) for each neuron, we ranked neurons by their relative contribution to frequency decoding (Fig. 2C). The contributions of individual PYRs to decoding were largely uniform across the population. This corroborates our earlier observation from the single-cell analysis that odor frequency discrimination does not predominantly occur at the level of individual neurons in the PCx.

Collectively, these analyses demonstrate that the odor frequency encoding is a robust capability of the PCx network and is highly distributed across the PYR population.

### Inhibition bidirectionally modulates stimulus encoding based on its mode of recruitment and task demands

In our PCx model, augmenting inhibition onto PYRs impaired odor frequency discrimination, whereas reducing it enhanced discrimination. This appears contrary to the general notion observed in OB and other cortical regions (59) that reducing inhibition impairs encoding capacity (72,73). Previous experimental studies have shown that enhancing inhibition of M/T cells in the mice OB improves odor discrimination, and vice versa (68–71,73,92).

This ambiguity partly stems from the mode of inhibition recruited and the specific demands of the discrimination task. For instance, disrupting centrifugal inhibition from the basal forebrain to OB granule cells causing an increased inhibition in the OB has been observed to produce a reversible impairment in olfactory discrimination (93). Furthermore, studies investigating targeted loss of Kv1.3 channels in M/Ts (94) and in the piriform cortex (95) reported enhanced odor discrimination associated with increased network excitability. Conversely, decreasing M/Ts excitability via activation of inhibitory DREADDs reduced odor discrimination (96). On the other hand, a study using a randomly connected network of cultured cortical neurons showed that blocking inhibition enhanced discrimination of stimuli originating from distal spatial loci, whereas the resolution for distinguishing adjacent inputs deteriorated (97). This finding demonstrated a spatially oriented task-dependent role of inhibition in stimulus discrimination by the cortical neurons. Such a dichotomy has been reported in several behavioral assays as well. For instance, GABA_A_ receptor β3-subunit-deficient mice outperformed wild-type mice in distinguishing closely related monomolecular alcohols but performed worse when distinguishing closely related mixtures of alcohols (69). Conversely, blocking GABAergic activity in the antennal lobe of honeybees has been shown to impair the discrimination of molecularly similar, but not dissimilar, odorants (66,67). In summary, the effect of inhibition on olfactory discrimination is not consistent but depends on the nature of the inhibitory mechanism recruited and the specific task demands.

Our findings suggest a trade-off between the network’s capacity to encode temporal dynamics and the dynamic scaling of inhibitory drive, a relationship that may be partly mediated by response gain, correlation, stochastic synchrony, and the underlying circuit architecture. In our study, eliminating either feedforward or feedback synaptic inhibition onto PYRs increased the response gain of pyramidal neurons. Response gain, a dynamic variable, facilitates adaptation of the network states to varying behavioral demands (98,99). Increased neural gain may produce robust encoding and optimize signal discrimination by facilitating attractor dynamics (100) and promoting a winner-take-all mechanism (101,102). For instance, different behavioral states during wakefulness such as arousal and locomotion are associated with increased gain and enhanced encoding of visual stimuli in the mouse primary visual cortex (103–105).

Global and recurrent inhibition in the olfactory circuit contribute to odor coding by suppressing noise and decorrelating neuronal responses. Substantial disruption of GABAergic inhibition is often associated with hypercorrelation, potentially driven in part by stochastic synchronization arising from shared noisy inputs in the absence of adequate inhibitory control. Although conventionally thought to degrade stimulus encoding, noise can paradoxically play a constructive role. Stochastic resonance has been shown to improve signal processing in both theoretical and experimental neuroscience models (106), and to enhance the reliability and regularity of neuronal firing across populations (107).

Every network, whether biological or simulated, has a unique optimal correlation structure. Thus, enhanced correlation may benefit neural coding depending on optimal correlation structure, stimulus statistics, population size, noise statistics, and decoding mechanisms employed (108–114).

This discrepancy in the role of inhibition may also be attributed to the stimulus selectivity of pyramidal neurons. In our PCx model, nearly all the pyramidal neurons were non-selective with respect to odor stimulus frequency (Fig. 1I–K). Accordingly, our study examined how inhibition shapes stimulus encoding in a non-selective neuronal population rather than addressing the more commonly studied case of selective neurons (35,74,115,116).

Further support for opposing directionality of inhibitory effects between OB and PCx arises from the difference in their cytoarchitecture and neuronal composition. In the OB, inhibitory neurons constitute approximately 80% and excitatory neurons approximately 20% of the total population, which is the inverse of the ratio observed in vertebrate cortex (117,118). Among these inhibitory neurons, granule cells are the predominant subtype (∼94%), inhibiting M/T cells via dense, reciprocal dendrodendritic synapses (119). This connectivity enables granule cells to exert strong lateral inhibition, positioning them as key regulators of odor tuning. Therefore, experimental perturbations that disrupted GABAergic inhibition in the OB by targeting granule cells resulted in impaired olfactory discrimination (68,69,72). In contrast, in our PCx model, inhibitory interneurons (feedforward and feedback subtypes) constitute approximately 17% of the neuronal population, while excitatory principal neurons account for the remaining ∼83%. More importantly, inhibition in the PCx is functionally specialized rather than globally dominant (41,120). Hence, the selective disruption of inhibition from individual interneuron subtypes, while constraining spontaneous/baseline firing of the network within physiological limits, may reconfigure circuit dynamics without abolishing inhibitory control altogether. Such reconfiguration may increase response separability, thereby improving decoding performance.

Notably, such effects likely reflect changes in population statistics rather than enhanced biological coding fidelity *per se*. Future *in vivo* and behavioral experiments will be required to validate our computational findings and resolve the apparent discrepancies. This will require the development of synapse-type-specific manipulations to generate targeted knockouts, as simulated in the present study. Nevertheless, our findings demonstrate that in a computational PCx model with non-selective pyramidal neurons, disrupting direct inhibition onto PYRs improves the discrimination of distantly related odor frequencies. Conversely, increasing inhibition impairs discrimination, highlighting the bidirectional capacity of the PCx circuit to modulate odor frequency encoding.

### Limitations of our model circuit

The current model simplifies PCx circuitry by not differentiating among various neuronal subtypes. We did not distinguish between different subclasses of principal neurons (e.g. semilunar cells vs. superficial pyramidal cells), feedforward interneurons (e.g., horizontal cells vs. neurogliaform cells) and feedback interneurons (e.g. regular-spiking multipolar vs. fast-spiking multipolar vs. bitufted vs. neurogliaform cells). Semilunar and superficial pyramidal neurons are functionally distinct subtypes due to their specialization in afferent and intracortical processing, respectively (42). Additionally, distinct interneuron subtypes contribute to layer-specific phasic inhibition of layer II principal neurons and hence play specific roles in cortical odor processing (41). In our model circuit we did not incorporate the centrifugal/feedback projections from the PCx back to the OB. These cortical feedback projections provide strongly normalized activity over diverse olfactory stimuli, which can be combined with the highly unbalanced activities of olfactory sensory neurons to influence M/T responses (29). The range of odor stimuli in our simulations was also limited in terms of mixture complexity, concentration, and frequency range. It would be intriguing to investigate the impact of these parameters on the frequency encoding capacity of the PCx. For instance, throughout our results, we observed poor decoding of odor frequency responses for stimulus Mix compared to stimuli 1 and 2.

Overall, our results suggest that the PCx network has the capacity to encode odor frequencies associated with naturalistic odor stimuli. Experimental validation of these findings would be valuable, as temporal dynamics in sensory stimuli are thought to be an important route of information transfer to the brain.

### Data availability

Source code for this study is openly available at https://doi.org/10.5281/zenodo.20270529

### Supplemental material

Supplemental Figures S1–S7: https://doi.org/10.5281/zenodo.20272407

## Acknowledgments

Preprint is available at https://doi.org/10.1101/2025.07.14.664633. This research utilized the High-Performance Cluster (Param Sanganak) facilities at the Computer Centre, IIT Kanpur. Graphical abstract created with BioRender.com.

## Disclaimers

The content is solely the responsibility of the authors and does not necessarily represent the official views of Indian Institute of Technology, Kanpur, 208016, India.

## Author Contributions

D.D. and A.K. conceived and designed the research. A.K. performed simulation experiments and analyzed data, prepared figures, and drafted the manuscript. A.K. and D.D. interpreted the results of the experiments, edited and revised the manuscript. A.K., O.C. & M.J. performed machine learning experiments and analyzed data with inputs from D.D. and A.M. The final version of the manuscript was approved by all the authors.

