## Supplemental Figures Legends for "Biophysically relevant network model of the piriform cortex predicts odor frequency encoding using network mechanisms"

**Figure S1.** Biophysical models of different piriform cortex neuron types. **A,** Schematic representation of a pyramidal neuron (PYR) model, consisting of a somatic compartment and five dendritic sections arranged from bottom to top. The model incorporates sodium (Na^+^), delayed rectifier potassium (K^+^), and leak ion channels, with their respective maximal conductance (gmax) and reversal potentials (E) indicated. **B,** Schematic of a single-compartment feedforward interneuron (FFIN) model. **C**, Schematic of a single-compartment feedback interneuron (FBIN) model, with the respective peak conductances and reversal potentials of the Na^+^, K^+^, and leak ion channels indicated.

**Figure S2.** Electrophysiological properties of a pyramidal neuron model. **A,** Step current injections and the corresponding voltage responses from the soma and five dendrites (Dend_sec1 to Dend_sec5). **B,** Measured steady-state voltage changes in the soma and dendritic sections as a function of injected current. The slope of linear regression for each trace gives the input resistance of the corresponding compartment. **C,** Action potentials evoked at suprathreshold current in each of the soma and dendritic sections. **D,** Mean spontaneous firing frequency (mean ± SD) recorded from each section across trials (*n* = 100). **E,** Plot of firing frequency vs. current step amplitude across all compartments. **F,** Example voltage vs. time traces of spontaneous activity recorded from the soma and each dendritic section.

**Figure S3.** Electrophysiological properties of a feedforward interneuron model (FFIN). **A,** Voltage responses recorded from the soma in response to three different step current injections (I to III). **B,** (I), Step current injections in the soma. (II), Corresponding voltage responses recorded from the soma. (III), Plot of change in voltage vs. injected current and its linear regression analysis to obtain the corresponding input resistance. **C,** Example action potential recorded from the soma (I) and the corresponding phase plot (II). **D,** Plot of firing frequency vs. current step amplitude. **E,** Example voltage traces of spontaneous activity recorded from the soma. **F,** Mean spontaneous firing frequency (mean ± SD) recorded across the trials (*n* = 100).

**Figure S4.** Electrophysiological properties of a feedback interneuron model (FBIN). **A,** Voltage responses recorded from the soma in response to step current injections (I to III). **B,** (I), Step current injections in the soma. (II), Corresponding voltage responses recorded from the soma. (III), Plot of change in voltage vs. injected current and its linear regression analysis to obtain the corresponding input resistance. **C,** Example action potential recorded from the soma (I) and the corresponding phase plot (II). **D,** Plot of firing frequency vs. current step amplitude. **E,** Example voltage traces of spontaneous activity recorded from the soma. **F,** Mean spontaneous firing frequency (mean ± SD) recorded across the trials (*n* = 100).

**Figure S5.** Decoding odor-frequency from baseline activity under various synaptic knockout conditions. Bar plots show 1D-CNNs’ classification accuracies for discriminating 2 Hz vs. 20 Hz odor conditions using only pre-stimulus/baseline (0–2 s) firing activity of all pyramidal neurons, alongside corresponding shuffle controls. Each panel represents a distinct synaptic disconnection: (**A**) Control, (**B**) without FF–PYR, (**C**) without FB–PYR, (**D**) without FF–FF, (**E**) without FB–FB, (**F**) without FF–FF and FB–FB, and (**G**) without PYR–PYR. Accuracy is reported across three stimulus conditions (stimulus 1, 2, and Mix), with each condition evaluated over 144 iterations using a random 60%/40% train-test split. Classification performance using baseline activity was statistically compared with chance-level decoding (shuffle control) using a two-tailed *t*-test (*p* < 0.05, *n* = 144).

**Figure S6.** Decoding accuracy as a function of pyramidal neuron (PYR) population size across control and knockout models using logistic regression classifiers. Mean classification accuracy for odor frequency discrimination (2 Hz vs. 20 Hz) for odor stimuli 1, 2, and Mix (left to right) is plotted against the number of neurons included in the analysis. Accuracy values were averaged across 144 iterations of random neuron subsampling (population step size = 25). Shaded regions represent ± SD. Chance-level performance was estimated by shuffling the trial labels. The logistic regression classifier was trained and tested for the following network configurations: without FF–PYR connections (W/o FF–PYR), without FB–PYR connections (W/o FB–PYR), without FF–FF connections (W/o FF–FF), without FB–FB connections (W/o FB–FB), without FF–FF & FB–FB connections (W/o FF–FF & FB–FB), and without PYR–PYR connections (W/o PYR–PYR) (top to bottom). The accuracy plot for the control network is shown in all panels for visual comparison.

**Figure S7.** Receiver operating characteristic (ROC) curves comparing the classification performance of 1D convolutional neural network models (1D-CNN; blue) and logistic regression classifiers (LRC; green) in decoding odor stimulation frequency (2 Hz vs. 20 Hz) for stimulus 1 (left), stimulus 2 (middle), and stimulus Mix (right) in control and knockout models arranged from top to bottom as follows: without FB–PYR connections (W/o FB–PYR), without FF–PYR connections (W/o FF–PYR), without FF–FF connections (W/o FF–FF), without FB–FB connections (W/o FB–FB), without FF–FF & FB–FB connections (W/o FF–FF & FB–FB), and without PYR–PYR connections (W/o PYR–PYR). The true positive rate (TPR) is plotted against the false positive rate (FPR) across decision thresholds (*n* = 144 iterations, pyramidal neuron population size = 500). The area under the ROC curve (AUC) quantifies overall decoding performance, with values near 0.5 indicating chance-level performance and values approaching 1.0 indicating perfect discrimination. AUC values are reported for CNNs and classifiers as mean ± SD (*n* = 144 for each). The dashed diagonal line denotes chance-level performance (AUC = 0.5).
